# Common molecular determinants underlie potyvirus host species jumps and resistance breakdown

**DOI:** 10.64898/2026.07.09.737469

**Authors:** Benoît Moury, Marion Szadkowski, Catherine Wipf-Scheibel C, Grégory Girardot, Julien Papaïx, Lionel Roques, Yamila Agrofoglio, Adrián A. Valli, Karine Berthier, Cécile Desbiez

**Affiliations:** INRAE, Pathologie Végétale, F-84140, Montfavet, France; INRAE, BioSP, 84914, Avignon, France; Centro Nacional de Biotecnología (CNB-CSIC), Madrid 28049, Spain

**Keywords:** Local adaptation, host adaptation, host shift, host range expansion, experimental evolution, VPg, *Potyvirus*, Asteraceae

## Abstract

Given their rapid evolutionary dynamics, viruses offer a powerful system to investigate the mechanisms underlying host jumps. Here, we experimentally evolved endive necrotic mosaic virus (ENMV) in five plant hosts within the family Asteraceae: two putative ancestral hosts (*Lactuca sativa* and *Tragopogon pratensis*), and three alternative crop or weed species (*Cichorium endivia*, *Zinnia elegans* and *Calendula arvensis*). The resulting evolved viral populations, together with the ancestral strain, were then evaluated in a reciprocal cross-inoculation experiment across all five host species.

ENMV exhibited clear adaptive responses in two hosts, *Z. elegans* and *C. arvensis*, with increased infection success and higher systemic viral accumulation compared to the ancestral virus. In contrast, no evidence of adaptation was detected in *L. sativa*, *T. pratensis* and *C. endivia*. Strikingly, strong cross-adaptation emerged between *Z. elegans* and *C. arvensis*: viral populations evolved in either host consistently outperformed those evolved in other hosts, as well as the ancestral strain, when infecting the reciprocal host.

Sequencing of the VPg cistron in adapted populations revealed multiple nonsynonymous mutations, several of which arose independently across evolutionary lineages and in both *Z. elegans* and *C. arvensis* selection regimes. Functional assays using an infectious ENMV cDNA clone demonstrated that seven of these substitutions, individually or in combination, significantly increased the infection rate in both *Z. elegans* and *C. arvensis*.

Notably, several of these substitutions also enhanced infectivity across four additional Asteraceae species among the eleven tested, without a clear relationship to host phylogenetic distance. Remarkably, all identified substitutions map to amino acid positions or adjacent residues in VPg previously implicated in the breakdown of recessive resistance genes against potyviruses in both crop and model plant systems. Together, these results suggest that adaptation to host resistance and host range expansion in potyviruses may rely, at least in part, on shared molecular pathways.

## Introduction

Parasite host range, *i.e.* the range of organisms where the parasite can carry out its infection cycle, is an important determinant of disease emergence. Indeed, Woolhouse and Gowtage-Sequeria (2005) have shown that the percentage of emerging or re-emerging human parasites increases with the extent of parasite’s host range. Over 40% of parasites with the widest host range (3 or more non-human host species) were emerging or re-emerging in human populations. This trend holds true for viruses, bacteria and fungi. Consistently, health crises are often linked to host jumps, the process by which parasites settle into a new host. Phylogenetic analyses provide evidence for recurrent host jumping (Moury and Desbiez 2020; Navaud et al. 2018) and the extinction of parasite lineages that did not jump to new hosts (Thines 2019).

In the case of plant viruses, jumps to new host genotypes within a given species are well documented. This process can be responsible for the breakdown of plant resistance and is usually determined by a small number (i.e. <3) of nonsynonymous substitutions in the virus genome (Gómez et al. 2009; Moury et al. 2021). Among more than 50 plant-virus combinations studied, only two cases do not follow this rule: the breakdown of melon resistance conferred by the *nsv* gene to *Melon necrotic spot virus* (MNSV, *Gammacarmovirus melonis*) (Truniger et al. 2008) and the breakdown of tomato resistance conferred by the *Ty-1* gene to begomoviruses (Belabess et al. 2015). In these cases, resistance breakdown involved recombination in a non-coding region of the virus genome. In the case of dominant monogenic plant resistance, almost all genomic regions of the virus can potentially be involved in resistance breakdown. In contrast, the majority of recessive monogenic resistances studied so far involve the VPg (viral protein genome-linked) coding-region as determinant of resistance breakdown (Tamisier et al. 2023, Moury et al. 2021).

Compared with intra-species virus jumps, virus adaptation or jump to new plant species is less well understood. On the basis of phylogenetic analyses, such trans-species jumps are considered to be fairly frequent. For example, Moury and Desbiez (2020) found that 3.1 host jumps per plant species occurred during the evolution of a set of 59 virus species belonging to the genus *Potyvirus*, a large group of plant viruses. The ecological factors and genetic events involved in host species jumps in plant parasites are largely unknown. Similarly, the plant genes involved in non-host resistance, i.e. the phenomenon whereby all members of a given plant species are resistant to all members of a given parasite species, are also largely unknown (Panstruga and Moscou 2020).

For plant viruses, only a few genetic determinants have been identified to induce host jump under natural epidemiological conditions. Most cases involve a few viral mutations, such as a mutation in the NIa-Pro cistron of papaya ringspot virus (PRSV, *Potyvirus papayanuli*) determining adaptation to papaya (Chen et al. 2008) and a mutation in the RNA-dependent RNA polymerase (NIb protein) involved in adaptation of plum pox virus (PPV, *Potyvirus plumpoxi*) to pea (*Pisum sativum*; Wallis et al. 2007). A few mutations in the P3 cistron are involved in adaptation of turnip mosaic virus (TuMV, *Potyvirus rapae*) to *Brassica* spp. or *Raphanus* spp. (Suehiro et al. 2004) and in adaptation of potato virus Y (PVY, *Potyvirus yituberosi*) to pepper (*Capsicum annuum*; Vassilakos et al. 2016). One or a few mutations in the VPg gene are involved in the adaptation of rice yellow mottle virus (RYMV, *Sobemovirus RYMV*) to the rice species *Oryza sativa* and *Oryza glaberrima* (Poulicard et al. 2012), potato virus A (PVA, *Potyvirus atuberosi*) to *Nicandra physalodes* (Rajamäki and Valkonen 1999), PPV to *Arabidopsis thaliana* and *Chenopodium foetidum* (Martinez-Turiño et al. 2021), tobacco etch virus (TEV, *Potyvirus nicotianainsculpentis*) to *A. thaliana* (Agudelo-Romero et al. 2008) and lettuce mosaic virus (LMV, *Potyvirus lactucae*) to *Catharanthus roseus* (Svanella-Dumas et al. 2014). The adaptation of citrus tristeza virus (CTV, *Closterovirus tristezae*) to different *Citrus* species is more complex, as it involves the acquisition of several genes by the virus (Tatiteni et al. 2011). Given the paucity of studies, many questions remain open about the genetic determinants and evolutionary mechanisms involved in host jumps by plant viruses.

We focused our study on endive necrotic mosaic virus (ENMV, *Potyvirus chichorii*), which belongs to the genus *Potyvirus* in the family Potyvirideae. ENMV is widespread in populations of wild salsify (*Tragopogon pratensis*) around Avignon in south-eastern France, where it reaches a prevalence of around 40% (Moury et al. unpublished results). It episodically infects lettuce crops, particularly American ‘Iceberg-type’ varieties, as most European lettuce cultivars have a high level of resistance to ENMV, probably linked to the *Tu* gene (Desbiez et al. 2017). The purpose of this study was to assess the risks of adaptation of ENMV to vegetable or ornamental crops, as well as to weed species that could serve as reservoirs. To this end, we carried out an experimental evolution of ENMV in several species of the family Asteraceae. We then identified mutations in the ENMV genome that were involved in host adaptation, and assessed the effect of these mutations on a wider set of plant species. This study highlighted the complexity of the effects of virus mutations on species host range, and the similarity of virus jumps between plant species with resistance breakdown at the intra-species level.

## Materials and Methods

### Plant and virus material

For the experimental evolution (EE), knowing that the host range of ENMV is restricted to the Asteraceae family (Desbiez et al. 2017), we chose five species belonging to three tribes of this family, namely Cichorieae, Calenduleae and Heliantheae: *Tragopogon pratensis* (Jack-go-to-bed-at-noon or meadow salsify, Cichorieae), *Lactuca sativa* cv. Calmar (lettuce, Cichorieae), *Cichorium endivia* var. latifolium cv. Géante maraîchère (escarole or endive, Cichorieae), *Calendula arvensis* (field marigold, Calenduleae) and *Zinnia elegans* (common zinnia, Heliantheae). *T. pratensis* is a biennial to perennial wild species widespread in the irrigated pastures surrounding the city of Avignon. *L. sativa* is a common vegetable crop, for which cultivars devoid of the *Tu* resistance gene are occasionally infected by ENMV. *C. endivia* and *Z. elegans* are vegetable and ornamental crops, respectively, for which natural ENMV infections have not been reported in France but that were shown to be susceptible to ENMV in the laboratory (Desbiez et al. 2017). Finally, *C. arvensis* is a wild plant not reported for natural ENMV infection that was shown to be partially susceptible to ENMV in the laboratory (Desbiez et al. 2017). We chose these species to test the potential of ENMV to adapt to various crop or wild Asteraceae species. Additional species of the tribes Cichorieae, Calenduleae and Heliantheae were later used for characterizing the host range of ENMV mutants.

We used ENMV isolate ENMV-FR (GenBank accession KU941946), collected from *L. sativa* cv. Montemar in Montfavet in November 2012 (Desbiez et al. 2017). In order to start the EE with a virus population as homogeneous as possible, transmission with a single *Myzus persicae* aphid was performed from a lettuce plant (cv. Calmar) mechanically inoculated with ENMV-FR. One inoculated lettuce plant (cv. Calmar) showing typical ENMV symptoms was used as a source for EE and for obtaining an infectious cDNA clone. This strain was named ENMV-7098MP1 and stored on the long term as dehydrated infected leaf material.

### Experimental evolution

Prior to the EE, we multiplied ENMV-7098MP1 by mechanical inoculation of *Cichorium endivia* var. latifolium cv. Géante maraîchère seedlings, in order to obtain high-titer inocula. For inoculation, virus-infected dehydrated leaves corresponding to one gram of fresh tissue was ground in 4 mL of phosphate buffer (0.03 M Na_2_HP0_4_, 0.2% sodium diethyldithiocarbamate), with 90 mg activated charcoal and 80 mg carborundum. Leaves from infected *C. endivia* plants were harvested 10 days post-inoculation (dpi) and used to prepare the inoculum for the first infection cycle of Asteraceae plants, which constituted the initial strain of the EE. For the EE (Fig. 1), we inoculated the first two leaves of 3-week-old seedlings (13 or 14 for *C. endivia*, *L. sativa*, *T. pratensis*, and *Z. elegans* and 98 for *C. arvensis* because its infection rate was expected to be low). Eight days later, the *C. arvensis* plants were inoculated a second time with the objective to increase the infection rate. Plants were arranged in a randomized block trial in a greenhouse. Twenty-six dpi, we harvested one gram of fully developed leaves from eight plants chosen randomly per plant species to initiate a second infection cycle for 40 evolutionary lineages (5 species x 8 independent replicates). For the second to sixth infection cycles, three 3-week-old seedlings were inoculated for each of the 40 evolutionary lineages (except for *C. arvensis* for which a higher number of seedlings was inoculated, 6 to 10, given its lower susceptibility). For each lineage, one of the infected plants was selected at random to initiate the next cycle, based on symptoms or ELISA detection.

**Figure 1.**
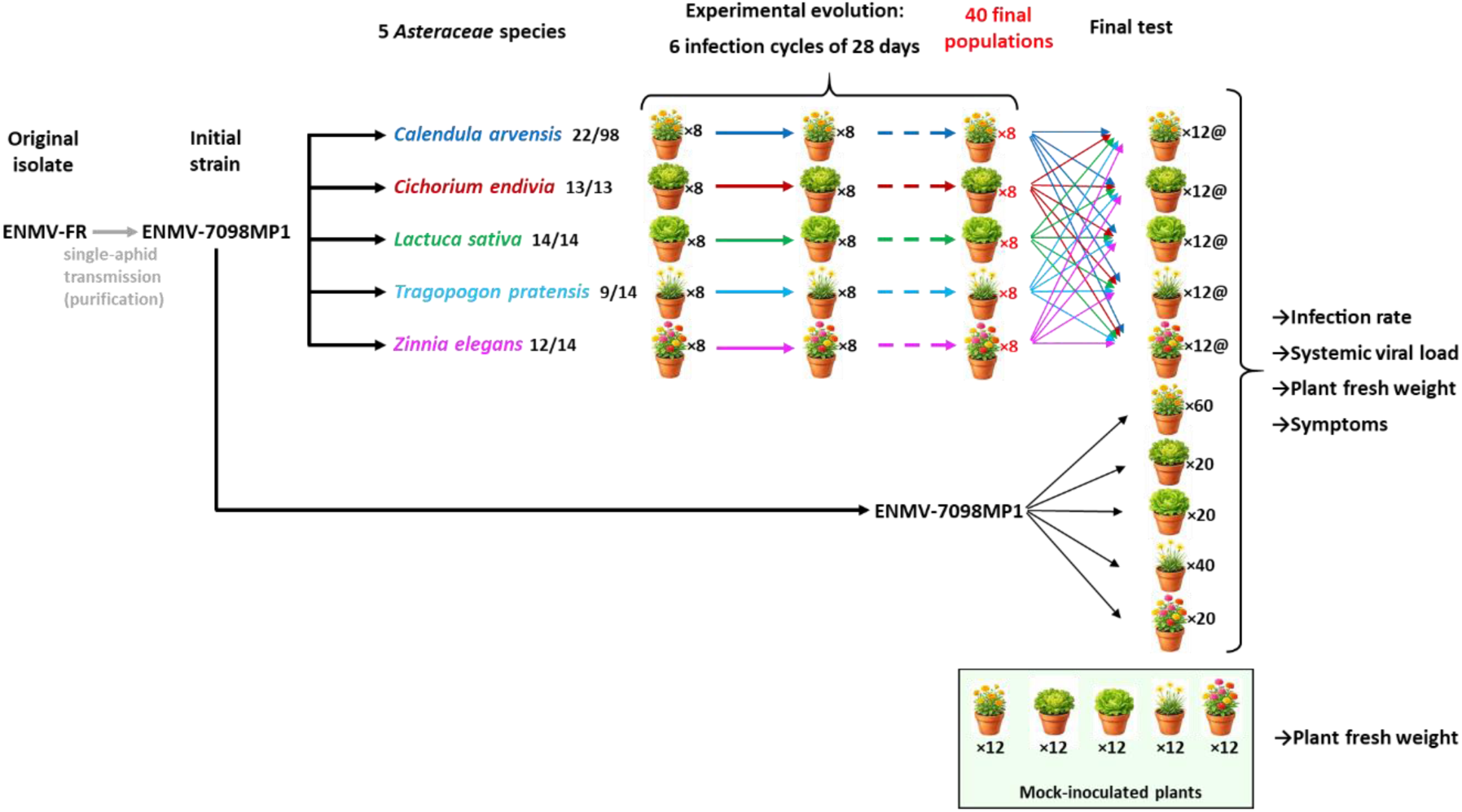
Experimental evolution (EE) design and phenotypic evaluation of ENMV initial strain and final populations. The EE consisted of six serial infection cycles on the same host species using ENMV-7098MP1 as initial inoculum source. Eight independent ENMV evolutionary lineages were obtained on each of five different Asteraceae species. After the EE, each of the 40 final ENMV populations was inoculated onto 12 plants of each of the five species and the initial strain ENMV-7098MP1 was inoculated onto 20 to 60 plants per species. The infection rate, systemic viral load, and fresh weight of infected plants were measured, and symptoms were recorded. The fresh weight of 12 mock-inoculated plants belonging to each plant species was also measured as a reference.

### Assessment of infectivity, systemic load and virulence of initial and final ENMV populations in a cross-inoculation assay

After the EE, we measured the infectivity, systemic load and virulence of the initial strain and final populations of ENMV on each of the five plant species (Fig. 1). Twelve seedlings were inoculated per final ENMV population and per plant species, using the plants infected at the end of the sixth infection cycle to produce inocula. The initial ENMV-7098MP1 strain was multiplied in *C. endivia* and inoculated to 20 to 60 plants, depending on the plant species (more plants of *C. arvensis* and *T. pratensis* were inoculated due to higher resistance or lower inoculation efficiency). In addition, we added 12 mock-inoculated seedlings per plant species. Plants were arranged in a randomized block trial in a greenhouse. Twenty-one dpi, we cut the plants at the cotyledon node and weighed them, and the systemic viral load was assessed by quantitative DAS-ELISA relative to a reference viral sample, as described in (Ayme et al., 2006).

### VPg sequencing of ENMV populations

One gram of apical leaf samples was ground in 3 mL extraction solution (Na_2_HPO_4_ 0.03M+0.2% diethyldithiocarbamate (DIECA)) in Bioreba® extraction bags. Two hundred microliters of extracts were used for RNA extraction with TRI-reagent (Molecular Research Center Inc., Cincinnati, OH) as described (Desbiez et al., 2009). Extracted RNAs were resuspended in 20 µL nuclease-free water and stored at - 20°C until use.

RT-PCR was performed with primers A0396 5’- CACACACTCGAGAACATCGC-3’ and A2112 5’-CCTTTTTGCAATGAGCGGGC-3’ yielding a 914-nt fragment encompassing the VPg-coding region. PCR products were sent to Genoscreen (Lille, France) for direct sequencing with both primers. Nucleotide sequences were aligned with MEGA6 (Tamura et al. 2013) and both nucleotide and deduced amino-acid sequences were compared to the original sequence of clone pLX-ENMV (see below).

### Building an infectious cDNA clone of ENMV

The pLX-ENMV clone (GenBank accession number PZ377050) was constructed by Gibson assembly from endive tissues infected with ENMV-7098MP1 (Agrofoglio et al., manuscript in preparation). The viral genomic cDNA sequence in this clone contains twenty nucleotide differences, seven of which corresponding to non-synonymous substitutions, compared to the reference sequence of ENMV-FR (GenBank accession number KU941946), which was obtained before single-aphid transmission.

In order to use yeast homologous recombination to build ENMV mutants, the vector of pLX-ENMV was replaced by a yeast-*Escherichia coli* binary vector. pLX-ENMV was digested with *Nco*I and *Bst*EII and a 12-kb fragment containing the whole ENMV sequence was purified from agarose gel and recombined in yeast YPH501 with plasmid pPRSV-E2 digested with *Eco*RI to remove most of the PRSV insert, as described in (Desbiez et al., 2012). DNA of yeast (*Saccharomyces cerevisiae* strain YHP501) growing on the selective CAU medium was extracted and used to transform electrocompetent *E. coli* DH5α. Bacterial plasmid DNA was extracted by alkaline lysis and controlled on 1% agarose gel after digestion with *Sal*I. The clone pENMV-11, presenting the expected restriction profile, was used for biolistic inoculation of susceptible *C. endivia* to control its infectivity (see below).

### Mutagenesis of the VPg cistron in pENMV-11

Eight single or double mutations were introduced in the VPg-coding region of the pENMV-11 clone by replacing the fragment between the two *Apa*LI sites at nucleotide positions 6650 and 7885 in the clone. Since several *Apa*LI sites were present in the vector, a yeast recombination strategy was used to assemble several overlapping DNA fragments (Supplementary Figure S1).

Two fragments of 8 and 3.5 kb resulting from the digestion of pENMV-11 with *Apa*LI were collected and purified from agarose gel (fragments 1 and 2). A third fragment of 7 kb resulting from the digestion of pENMV-11 with *Sal*I was also purified (fragment 3). The VPg-coding fragments of seven natural variants (Table 1) were amplified with primers A0396 and A2112 (Supplementary Table S1) using a high-fidelity DNA polymerase. Another fragment was amplified to fill the gap between the position of primer A2112 and the *Apa*LI site at position 7886 in the intron-containing clone, and fusion PCR was used as described in (Desbiez et al., 2012) to bind these two fragments, yielding fragment 4 that contained the desired mutation(s). In addition, since no natural isolate was found to possess mutation H122N only, this mutant was obtained by site-directed mutagenesis as described in (Desbiez et al., 2014) using mutagenic primers A2129-A2130, and A2128-A2097 as external primers (Supplementary Table S1).

**Table 1.** Non-synonymous substitutions observed in the VPg cistron of the final ENMV populations.

| ENMV | Amino acid position |  |  |  |  |  |  |  |  |  |  |
| --- | --- | --- | --- | --- | --- | --- | --- | --- | --- | --- | --- |
|  | 23 | 56 | 94 | 104 | 122 | 123 | 124 | 125 | 165 | 183 | 185 |
| Wild-type | F | M | V | E | H | N | T | I | G | E | E |
| Final populations from *C. arvensis* |  |  |  |  |  |  |  |  |  |  |  |
| 1<sup>a</sup> |  |  |  |  | R | T |  |  |  |  |  |
| 2<sup>a</sup> |  |  |  |  |  |  |  |  | C |  |  |
| 3<sup>a</sup> |  |  |  |  | R |  |  |  | S |  |  |
| 4 |  |  |  |  | R |  |  |  | S |  | K |
| 5 |  |  |  |  | R |  |  |  | S | K |  |
| 6 |  |  |  | E/V<sup>b</sup> | R | N/T<sup>b</sup> |  |  | G/S<sup>b</sup> |  |  |
| 7 |  |  |  |  |  |  |  |  | C |  |  |
| 8 |  |  |  |  |  |  |  |  | C |  |  |
| Final populations from *Z. elegans* |  |  |  |  |  |  |  |  |  |  |  |
| 1<sup>a</sup> |  |  |  |  | N |  |  | V |  |  |  |
| 2 | C |  |  |  |  |  |  | L |  |  |  |
| 3 |  |  |  |  |  |  |  |  | C |  |  |
| 4<sup>a</sup> |  |  |  |  | R |  |  |  |  |  |  |
| 5<sup>a</sup> |  |  |  |  |  |  |  | L |  |  |  |
| 6 |  | T |  |  |  |  |  | L |  |  |  |
| 7 |  |  | I |  |  |  |  | L |  |  |  |
| 8<sup>a</sup> |  |  |  |  | N |  | K |  |  |  |  |

<sup>a</sup>: ENMV populations used to amplify the VPg cistron and to build mutant clones.
<sup>b</sup>: at the given position, two nucleotides corresponding to two amino acids are in mixture.

For each mutant, yeast recombination was performed by mixing plasmid fragments 1 to 4. After growth on selective CAU medium, yeast plasmid DNA was extracted and used to transform electrocompetent *E. coli* DH5α. Bacterial plasmid DNA was extracted by alkaline lysis and controlled on 1% agarose gel after digestion with *Xho*I or *Pvu*II.

### Biolistic inoculation of infectious clones

Biolistic inoculation of infectious clones was performed as described by Romay et al. (2015). Three microlitres of plasmid DNA were mixed with 3 µL H_2_O, 30 µL of a tungsten suspension (50 mg.mL^-1^ in 50% glycerol) and 30 µL 2.5M CaCl_2_. After 5 min on ice, the mixture was centrifuged for 3 mn at maximum speed in a microcentrifuge, the supernatant was removed carefully and 100 µL 70° ethanol were added. After a second centrifugation for 3 min at maximum speed, the supernatant was removed and 100 µL of absolute ethanol were added. Eight microlitres were used for each plant bombardment using a biolistic delivery system (Gal-On et al., 1997) with 4 bar compressed air. The bombardments were performed on young leaves of 4-week-old *C. endivia* plantlets, and the plants were grown in the greenhouse until symptom development. For one symptomatic plant for each mutant, RNA was extracted with TRI-reagent and the presence of the expected mutations was controlled by partial Sanger sequencing with primer A0396 as described above.

### Host range evaluation of ENMV VPg mutants

To test if VPg mutations involved in adaptation of ENMV-FR to *C. arvensis* or *Z. elegans* resulted in broader adaptation to particular tribes of the Asteraceae, we inoculated the wild-type ENMV-FR and seven VPg mutants on 14 species belonging to the three tribes corresponding to the species used for EE. The results obtained with the eighth VPg mutant could not be interpreted, either because its progeny contained an additional nonsynonymous mutation in the VPg, or because the inoculum concentration was very low (see Results). These plant species belonged to the tribes Calenduleae (*C. arvensis*, *C. officinalis*, *Dimorphotheca pluvialis* cv. Polar star and *Osteospermum ecklonis* cv. F_1_ Passion mixed), Heliantheae (*Z. elegans*, *Cosmos bipinnatus* cv. Virgo, *Rudbeckia hirta* cv. Marmalade, *Helianthus annuus* cv. Sunny smile, *Echinacea purpurea* cv. Magnus and *Coreopsis grandiflora* cv. Early sunrise) and Cichorieae (*C. endivia*, *Catananche caerulea*, *Helminthotheca echioides* and *Urospermum dalechampii*). Seeds were from Graines Baumaux (Mazirot, France) or Voltz Horticulture (Loire-Authion, France), except those of *H. echioides* and *U. dalechampii*, which we harvested on the site of the Pathologie Végétale unit in Montfavet.

We used *C. endivia* plants inoculated by biolistics as inoculum sources to inoculate 10 to 60 plants per species and per virus, depending on the plant susceptibility, with inocula calibrated by quantitative DAS-ELISA. Infection was evaluated by symptom assessment and ELISA. For *C. arvensis* and *Z. elegans*, we performed quantitative DAS-ELISAs to compare the systemic load of ENMV-FR and its mutants, as described in (Ayme et al. 2006). The sequence of the VPg-coding region of the viral progeny was determined for 1 to 5 randomly chosen infected plants for each plant species and each ENMV variant, as described above.

## Statistical analyses

We performed all statistical analyses using R software version 4.4.3. The R Markdown script covering all the statistical analyses performed in this article and the datasets are available online at https://doi.org/10.5281/zenodo.21099388. The versions of the R packages used are indicated in Supplementary Table S2.

For analysis of infection rates of the cross-inoculation assay, we used a Bayesian binomial mixed-effects model fitted with the package *brms* (Bürkner 2017, 2018). *brms* relies on Stan probabilistic language, which implements Bayesian inference through Markov chain Monte Carlo methods (Carpenter et al., 2017). The model included the ‘test species’, the ‘viral population type’ (both initial strain and evolved populations) and their interaction as fixed effects, and took into account the ‘experimental block’ and ‘virus lineage’ (nested within the ‘viral population type’) as random effects. The Markov chains were run with 4 chains, each with 4,000 iterations of which the first 2,000 were warmup to calibrate the sampler, leading to a total of 8,000 posterior samples. To assess the robustness of our results to prior assumptions, we performed a sensitivity analysis of the fixed-effect priors, comparing Normal distributions with standard deviations of 1, 2, and 5 (***N***(0,1), ***N***(0,2) and ***N***(0,5)). Models were evaluated using leave-one-out cross-validation with the *loo* package (Vehtari et al. 2017).

For a given test species, we also tested whether the infection rates of ENMV populations evolved on the different plant species differed significantly from those of the initial strain using Fisher’s exact tests. Fisher’s exact tests were also used to compare the infection rates of the wild-type ENMV clone and its VPg mutants.

For analysis of systemic viral loads and plant fresh weights of the cross-inoculation assay, we used linear mixed-effects models implemented in the *lme4* package, where the ‘test species’ and ‘viral population type’ (i.e. either the initial ENMV strain or ENMV populations that had evolved in each of the five plant species) were fixed effects, ‘experimental block’ was a random effect, and ‘virus lineage’ was a random effect nested within the ‘viral population type’ effect. For both traits, residual normality and homoscedasticity were assessed both graphically and using the *DHARMa* package.

Local adaptation in plant hosts during the experimental evolution was assessed using complementary foreign-versus-local and home-versus-away frameworks (Kawecki and Ebert 2004). In the foreign-versus-local framework, *local* viral populations are those that were experimentally evolved on the same host species as the focal plant (i.e. sympatric combinations), whereas *foreign* viral populations are those evolved on alternative host species (i.e. allopatric combinations). Viral performance, i.e. infection rate and systemic viral load, was therefore compared between locally and foreign-evolved viral populations for each plant species. In the home-versus-away framework, *home* refers to the host species on which a given viral population evolved, while *away* refers to alternative host species used in the assay, allowing comparison of viral performance in home versus away hosts within each viral population type.

For these analyses, we used models similar to those described above, but incorporating framework-specific fixed effects. In the foreign-versus-local analyses, models included ‘test species’, a binary ‘local’ variable (local vs. foreign viral populations), and their interaction. In the home-versus-away analyses, models included ‘viral population type’, a binary ‘home’ variable (home vs. away hosts), and their interaction. Data from the ancestral strain were excluded from all analyses.

Finally, systemic loads of the wild-type ENMV clone and its VPg mutants in *C. arvensis* and *Z. elegans* were compared using analysis of variance (ANOVA) followed by Tukey HSD post hoc tests. Viral load values in *C. arvensis* were Box–Cox transformed prior to analysis to satisfy normality of residuals and homogeneity of variances. A linear model (function *lm*) was fitted including ‘ENMV variant’ and ‘experimental block’ as fixed effects. The interaction between ENMV variant and block was tested and found to be non-significant; therefore, it was excluded from the final model. Pairwise comparisons among ENMV variants were conducted using Tukey-adjusted post hoc tests based on estimated marginal means with the package *emmeans*. Viral load values in *Z. elegans* were log transformed prior to analysis to satisfy normality and homogeneity of variances. The interaction between ENMV variant and block was found to be significant; therefore, the statistical analysis was conducted separately for each block.

## Results

### Adaptation of ENMV to *Calendula arvensis* and *Zinnia elegans* following experimental evolution

According to DAS-ELISA detection, the initial ENMV-7098MP1 strain infected a low proportion (22.4%) of *C. arvensis* plants during the first infection cycle, whereas higher infection percentages (64.3% and 85.7%) were observed in *T. pratensis* and *Z. elegans*, respectively, and infection rates of 100% were observed in *C. endivia* and *L. sativa* (Fig. 2). During the experimental evolution of ENMV, 100% of *C. arvensis* plants were infected by the third infection cycle, and this percentage remained stable until the end of the experiment. Similarly, 100% of *Z. elegans* plants were infected by the end of the fourth infection cycle, a rate that remained stable thereafter. In contrast, the infection rate of *T. pratensis* fluctuated over the course of the experiment (41.7 to 60.7%) but did not increase significantly. Finally, the infection rate remained stable at 100% in *C. endivia* and *L. sativa*. These results suggest an adaptation of ENMV in *C. arvensis* and possibly *Z. elegans* during experimental evolution. The rather low percentage of infection in *T. pratensis* may be due either to the filiform shape of its leaves, which are not easy to inoculate mechanically and may induce infection escapes, or to a heterogeneity of resistance of the plants, since the seeds were collected from a natural local population.

**Figure 2.**
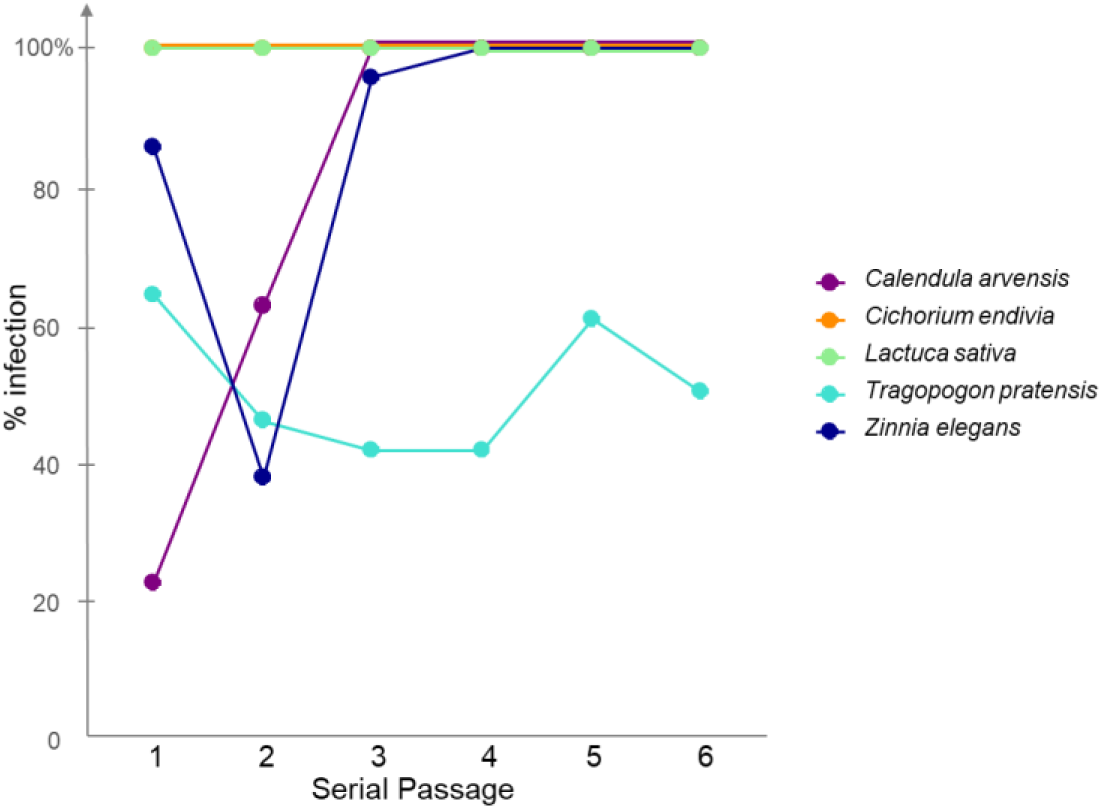
Systemic infection rate by ENMV in the five plant species used for experimental evolution during six consecutive infection cycles.

The symptoms induced by ENMV differed depending on the host plant species. In *L. sativa* and *C. endivia*, severe mosaic patterns were observed on the leaves, and infected plants showed stunted growth, regardless of the viral strain or population. In contrast, symptoms of streak mosaic were barely visible on the leaves of infected *T. pratensis* plants. Finally, infected *C. arvensis* and *Z. elegans* plants showed mild mosaic and a mixture of mosaic and necrosis on the apical leaves, respectively.

After the six infection cycles, all evolved ENMV populations and the initial strain were inoculated to the five host species in a cross-inoculation assay. In this assay, for the inoculated *C. arvensis* plants, no infection was observed for the initial strain or for the eight populations evolved in *C. endivia* (Fig. 3A). Similarly, the infection rates were low for the populations evolved in *L. sativa* or in *T. pratensis* (5.2% and 2.1% on average, respectively). In contrast, infection rates were very high for the populations evolved in either *C. arvensis* or *Z. elegans* (100% and 85.4% on average, respectively), except for one of the eight ENMV populations evolved in *Z. elegans* that was not infectious in *C. arvensis*. Similar results were observed for inoculated *Z. elegans* plants. Low infection percentages were observed for the initial strain and for the populations evolved in *C. endivia*, *L. sativa* or *T. pratensis* (20.0%, 15.6%, 16.7% and 14.6% on average, respectively). In contrast, high infection rates were observed for the populations evolved in *C. arvensis* or *Z. elegans* (80.2% and 92.7% on average, respectively).

**Figure 3.**
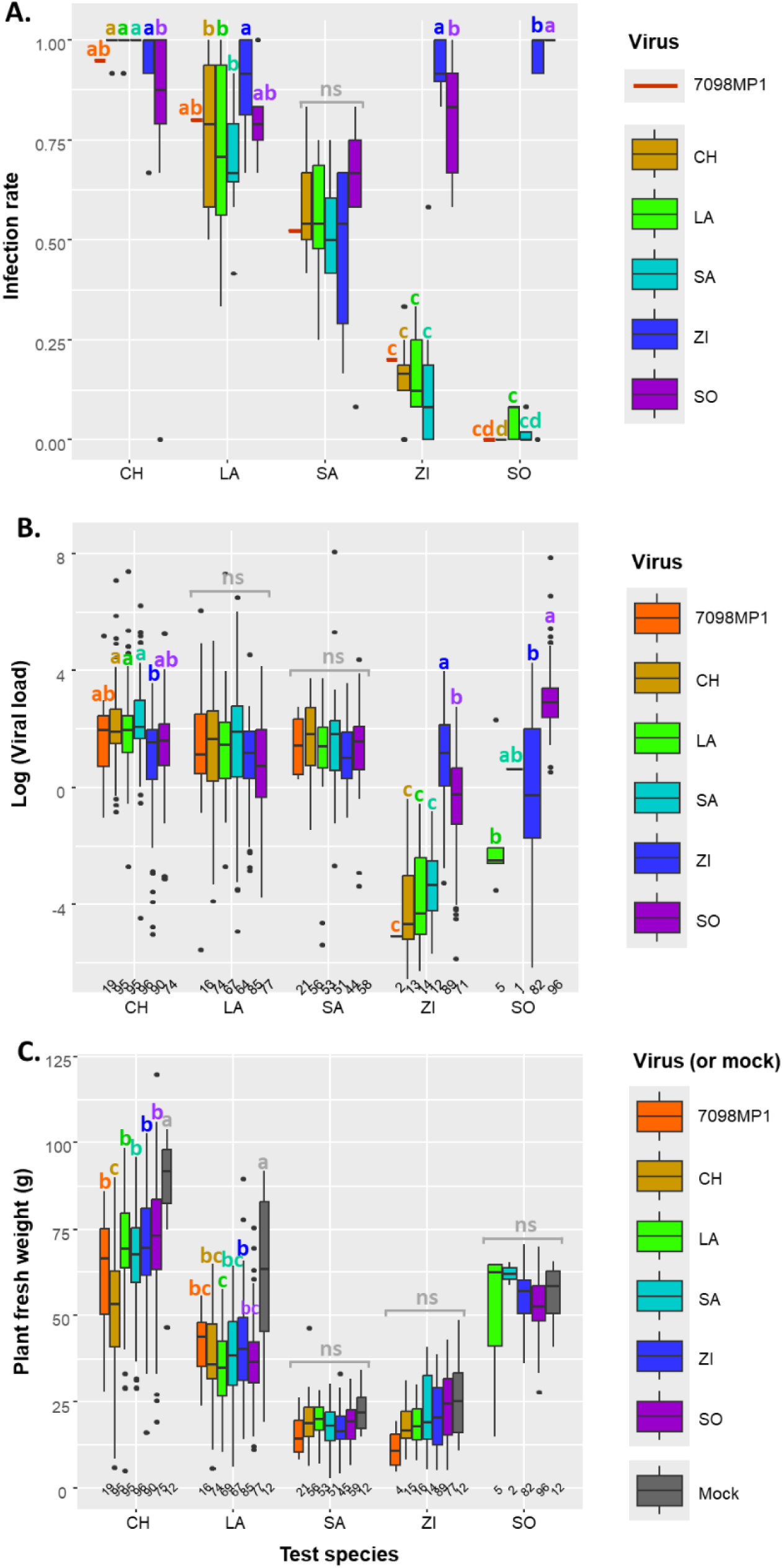
Phenotypes of the final ENMV populations and the initial strain ENMV-7098MP1 on the five plant species used for experimental evolution, as observed from the cross-inoculation assay. **A.** Infection rate. The Bayesian binomial mixed-effects model used for statistical analysis (brm function) was ‘infection rate’ ∼ ‘test species’ * ‘viral population type’ + (1|‘block’) + (1|‘viral population type’:‘lineage’). **B.** Systemic viral load. The linear mixed-effects model used for statistical analysis (lmer function) was ‘log(systemic viral load)’ ∼ ‘test species’ * ‘viral population type’ + (1|‘block’) + (1|‘viral population type’:‘lineage’). **C.** Plant fresh weight. The mixed model used for statistical analysis (lmer function) was ‘fresh weight’ ∼ ‘test species’ * ‘viral population type’ + (1|‘block’) + (1|‘viral population type’:‘lineage’). Boxes indicate the interquartile range (IQR), horizontal lines the medians, and whiskers extend to the most extreme data points that are within 1.5 × IQR from the box.

We analysed infection rates using a Bayesian binomial mixed-effects model. A sensitivity analysis of the priors for fixed effects (***N***(0,1), ***N***(0,2), ***N***(0,5)) showed that posterior means were robust, with extensive overlap across priors. Wider 95% credible intervals for ***N***(0,5) reflected increased uncertainty under a more permissive prior, whereas intervals were narrower for ***N***(0,2) and ***N***(0,1). Leave-one-out cross-validation (LOO) indicated that ***N***(0,2) maximized predictive accuracy (expected log pointwise predictive density; Vehtari et al., 2017) while minimizing influential observations (Pareto k), whereas ***N***(0,1) was slightly restrictive and ***N***(0,5) produced more influential points. Based on predictive performance and influence diagnostics, ***N***(0,2) was chosen for the final model.

The 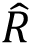 values were equal to 1.00, indicating that convergence of the sampling distribution to a stationary distribution has likely occurred (Gelman and Rubin 1992). Effective sample sizes were well above recommended thresholds (bulk and tail ESS > 3100; Johnston et al. 2024). No divergent transitions were detected, indicating robust posterior exploration. Posterior predictive checks confirmed close agreement between observed and simulated outcomes, including extreme binomial events. Residual overdispersion was low (1.21), supporting adequate model fit. Overall, predictive performance was robust, and model inferences are reliable.

Random-effect estimates indicated moderate variation among experimental blocks (standard deviation = 0.55; 95% credibility interval: [0.16, 1.70]) and substantial variation among lineages nested within virus population type (standard deviation = 0.57; 95% credibility interval: [0.39, 0.80]), reflecting biologically meaningful heterogeneity among lineages.

Any points beyond the whiskers are plotted individually as outliers. Letters correspond to significant differences between virus treatments for a given test species (ns: not significant).

CH: *Cichorium endivia*; LA: *Lactuca sativa*; SA: *Tragopogon pratensis*; ZI: *Zinnia elegans*; SO: *Calendula arvensis*.

Mock: mock-inoculated plants.

The numbers of infected plants or mock-inoculated plants in each category are indicated at the bottom of panels B and C.

The model revealed a strong interaction between test species and virus population type (Supplementary Fig. S2). Infection rates varied among species for a given viral population type, following a gradient from susceptibility to resistance (*C. endivia* > *L. sativa* > *T. pratensis* > *Z. elegans* > *C. arvensis*). This gradient did not apply for populations evolved in *C. arvensis* or *Z. elegans*. In these cases, infection rates in *T. pratensis* were significantly lower than in other species. Populations evolved in *C. arvensis* also showed lower infection in *T. pratensis*, and infection rates in *C. endivia*, *L. sativa*, and *Z. elegans* were lower than in *C. arvensis*. Comparison between virus population types revealed no or only few significant differences in *C. endivia*, *L. sativa*, or *T. pratensis* (Fig. 3A). In contrast, populations evolved in *C. arvensis* or *Z. elegans* exhibited significantly higher infection rates than other populations, including the initial strain, in both *C. arvensis* and *Z. elegans*. These results suggest cross-adaptation of ENMV-7098MP1 in *Z. elegans* and *C. arvensis* during experimental evolution. Fisher’s exact tests confirmed that only ENMV populations evolved in *C. arvensis* or *Z. elegans* showed a significant increase in infection rates compared with the initial ENMV-7098MP1 strain (p-values < 0.027), with the exception of a single population evolved in *Z. elegans* that did not infect *C. arvensis*.

To analyse systemic viral loads and plant fresh weights, we used linear mixed-effects models with the ‘test species’, ‘viral population type’ and their interaction as fixed effects and taking into account the ‘experimental block’ and ‘virus lineage’ as random effects. After logarithmic transformation for viral loads, diagnostics performed with the *DHARMa* package revealed a few outliers and a slight deviation from uniformity in the residual distribution (Kolmogorov-Smirnov test, p < 0.01), but no overdispersion was detected, and the residuals did not show any systematic pattern or curvature. For plant fresh weights, conclusions regarding the fixed effects remained largely unchanged after logarithmic or square-root transformations, indicating that the model was robust despite these minor deviations. Analysis of systemic viral loads revealed a significant interaction between the effects of the ‘test species’ and the ‘viral population type’ (Fig. 3B). In *L. sativa* or *T. pratensis*, no significant differences were observed between virus population types (initial ENMV strain or ENMV populations that had evolved in each of the five plant species). In *C. endivia*, the systemic loads of ENMV populations evolved in *C. endivia*, *L. sativa* or *T. pratensis* were significantly higher than that of populations evolved in *Z. elegans*, while the populations evolved in *C. arvensis* and the initial strain did not show any significant difference compared to the others and between each other. In *C. arvensis* infected plants, significantly higher systemic viral loads were observed for virus populations evolved in the same species (*C. arvensis*) than for virus populations evolved in *Z. elegans* or *L. sativa*. A single *C. arvensis* plant was infected by ENMV populations evolved in *T. pratensis*. Finally, with regard to virus loads in infected *Z. elegans* plants, three categories of viral origins could be distinguished. Viral populations that evolved within the same species (*Z. elegans*) had significantly higher systemic loads than populations that evolved in *C. arvensis*, which in turn, had significantly higher systemic loads than other evolved populations or the initial strain. The viral load data confirmed that ENMV adapted to *C. arvensis* and *Z. elegans* during the EE, and that cross-adaptation occurred between these two host species. However, this cross-adaptation was incomplete, as viral loads were consistently higher in « local » combinations (i.e., ENMV populations infecting the same host species in which they had evolved) than in « foreign » combinations (i.e., ENMV populations evolved in *C. arvensis* and infecting *Z. elegans*, or vice versa).

With regard to the fresh weight of plants, the linear mixed-effects model also revealed a significant interaction between the effects of the ‘test species’ and those of the ‘virus population type’ (Fig. 3C). In *T. pratensis*, *Z. elegans* and *C. arvensis*, no significant differences were observed between mock-inoculated plants, plants infected with the initial ENMV strain, and plants infected with evolved populations. This suggests a high level of tolerance of these plant species. In contrast, in *C. endivia* and *L. sativa*, mock-inoculated plants had a significantly higher fresh weight than infected plants, suggesting a lower tolerance of these plant species. For these two plant species, the only significant differences between the viruses was (i) that the fresh weight of *C. endivia* plants was lower when they were infected with ENMV populations evolved in *C. endivia* than when they were infected with other evolved ENMV populations or with the initial strain and (ii) that the fresh weight of *L. sativa* plants was lower when they were infected with populations evolved in *L. sativa* than when they were infected with populations evolved in *Z. elegans*. This suggests an increase in ENMV virulence during EE in *C. endivia* or *L. sativa* in these respective host species compared to EE in other plant species. These changes in virulence were not associated with significant changes in viral load.

Using the local-versus-foreign framework, we detected strong local adaptation for infection rate and systemic viral load in *Z. elegans* and *C. arvensis*, with consistently higher performance of local than foreign viral populations. A weaker but significant signal was observed in *C. endivia*, whereas no evidence of local adaptation was found in *L. sativa* or *T. pratensis* (Supplementary Fig. S3). The home-versus-away framework yielded consistent patterns, with several viral population types showing higher performance in their home host than in away hosts. This effect was strongest for viral populations evolved on *C. endivia* and *C. arvensis*, intermediate for those evolved on *Z. elegans*, and absent for those evolved on *T. pratensis*. Viral populations evolved on *L. sativa* showed higher infection probability in their home host but no difference in systemic viral load relative to away hosts.

In summary, these results demonstrate the adaptation of ENMV in *C. arvensis* and *Z. elegans*, as well as cross-adaptation of ENMV between these two host species. These adaptations did not result in any detectable fitness costs in the host species *L. sativa* or *T. pratensis*. In *C. endivia* however, a significant cost in terms of infection rate was observed for viral populations evolved in *C. arvensis*, compared to populations evolved locally (i.e. in *C. endivia*) (Fig. 3A). Likewise, a significant cost in terms of systemic viral load was detected for populations evolved in *Z. elegans*, compared to locally-evolved populations (Fig. 3B). Viral evolution also had only a limited effect on its impact on plant fresh weight. Nevertheless, increases in virulence (i.e. reductions in fresh weight) were observed for virus populations evolved on *C. endivia* and *L. sativa* when infecting their respective local host species, compared to virus populations evolved on other species (Fig. 3C).

### Mutations in the VPg cistron are responsible for ENMV adaptation to new host species

We analysed the genome of evolved ENMV populations, sampled at the end of the sixth infection cycle, to identify candidate mutations for host adaptation. To do this, we first focused on the VPg cistron, which has been shown to be involved in the adaptation of several potyviruses to host plant immunity or in virus host shifts (Rajamäki and Valkonen 1999, Agudelo-Romero et al 2008, Svanella-Dumas et al 2014, Martinez-Turiño et al 2021, Tamisier et al 2023). The 16 ENMV populations evolved in *C. arvensis* or *Z. elegans* presented the strongest evidence for adaptation. From one to four nonsynonymous substitutions were observed in the VPg cistron of these populations (Table 1). In addition, parallel mutations were observed among the eight experimental replicates at codon positions 122, 123 and 165 for *C. arvensis*, and at codon positions 122 and 125 for *Z. elegans*, strongly confirming their adaptive role. Finally, the same mutations at codon positions 122 and 165 were observed for ENMV populations evolved in *C. arvensis* and *Z. elegans*, providing a possible explanation to the cross-adaptations observed in these two plant species (Fig. 3).

In contrast, no nonsynonymous substitutions were observed in ENMV populations evolved in *C. endivia*, *L. sativa* or *T. pratensis* (3 randomly selected populations were sequenced for each plant species). Of these 9 populations, only one non-fixed synonymous substitution was detected in a population evolved in *C. endivia*.

We used a reverse genetics approach to analyze the involvement of some of the VPg mutations in ENMV adaptation. Using an infectious cDNA clone derived from the initial strain used for experimental evolution, we constructed eight single or double VPg mutants. We focused on the most frequently detected mutations, at positions 122, 123, 125 and 165. The H122N mutation was only detected in combination with another (T124K or I125V). Consequently, we constructed these two double mutants in addition to the single H122N mutant to test for a cumulative effect of mutations on the level of adaptation of ENMV. We used one of the most susceptible hosts (*C. endivia*) to perform biolistic inoculations with the mutant cDNA clones to obtain high titer inocula. In two independent experiments, the G165C mutant either accumulated at a low titer in *C. endivia* or had an additional unfixed VPg mutation (T120I). Consequently, it was not possible to analyze the effect of the G165C mutation alone on the level of adaptation of ENMV.

The remaining seven mutants infected *C. arvensis* at a significantly higher rate (30.0 to 85.0%) than the wild-type clone (1.7%) (Table 2). Similar results were observed in *Z. elegans*, where the seven mutants infected from 70.0 to 100.0% of inoculated plants, compared with 5.0% of plants infected by the wild-type clone.

**Table 2.** Systemic infectivity of ENMV variants in eight Asteraceae species. For a given plant species, infection rates were compared with those of wild-type ENMV using Fisher’s exact tests and Bonferroni corrections. *: p<0.05; **: p<0.01; ***: p<0.001.

| ENMV<br>variant | Tribe Calenduleae |  |  | Tribe Heliantheae |  |  | Tribe Cichorieae |  |
| --- | --- | --- | --- | --- | --- | --- | --- | --- |
|  | *C. arvensis* | *D. pluvialis* | *O. ecklonis* | *Z. elegans* | *C. bipinnatus* | *R. hirta* | *U. dalechampii* | *H. echinoides* |
| Wild type | 1/60 (1.7)<sup>a</sup> | 43/50 (86.0) | 0/41 (0.0) | 3/60 (5.0) | 0/60 (0.0) | 3/45 (6.7) | 13/49 (26.5) | 9/20 (45.0) |
| 122N | 18/60 (30.0)*** | 18/20 (90.0) | 0/19 (0.0) | 40/40 (100.0)*** | 2/20 (10.0) | 1/19 (5.3) | 17/20 (85.0)*** | 3/10 (30.0) |
| 122N+124K | 26/40 (65.0)*** | 19/20 (95.0) | 0/20 (0.0) | 39/40 (97.5)*** | 8/20 (40.0)*** | 1/18 (5.6) | 34/40 (85.0)*** | 4/10 (40.0) |
| 122N+125V | 21/59 (35.6)*** | 19/20 (95.0) | 0/20 (0.0) | 40/40 (100.0)*** | 2/20 (10.0) | 6/39 (15.4) | 24/40 (60.0)* | 4/10 (40.0) |
| 122R | 34/40 (85.0)*** | 20/20 (100.0) | 0/20 (0.0) | 28/40 (70.0)*** | 19/40 (47.5)*** | 17/39 (43.6)*** | 11/20 (55.0) | 5/10 (50.0) |
| 122R+123T | 31/40 (77.5)*** | 20/20 (100.0) | 6/28* (21.4) | 32/40 (80.0)*** | 7/40 (17.5)** | 21/32 (65.6)*** | 36/40 (90.0)*** | 3/10 (30.0) |
| 122R+165S | 31/40 (77.5)*** | 12/20 (60.0) | 0/20 (0.0) | 38/40 (95.0)*** | 0/20 (0.0) | 8/38 (21.1) | 9/19 (47.4) | 4/10 (40.0) |
| 125L | 15/40 (37.5)*** | 13/40 (32.5)*** | 0/20 (0.0) | 39/40 (97.5)*** | 2/20 (10.0) | 2/18 (11.1) | 4/20 (20.0) | 3/10 (30.0) |

<sup>a</sup> Number of infected plants / total number of inoculated plants. Percentages of infected plants are in brackets.

In *Z. elegans*-infected plants, the VPg sequence of mutant progeny matched that of the corresponding cDNA clone in the vast majority of cases (21/22) (Table 3). The only exception was one of the four progenies of the 122N mutant, which fixed the additional 165C mutation. In contrast, the two infected plants inoculated with wild-type ENMV that we analysed (out of 3 in total) showed fixation of the additional 125L mutation, which has already been detected in four of the eight ENMV populations evolved in *Z. elegans* (Table 3). These results indicate that the seven mutations or mutation combinations we analysed with the mutant cDNA clones were sufficient to confer a high level of adaptation in *Z. elegans* to ENMV, consistent with their parallel fixation during experimental evolution.

**Table 3.**
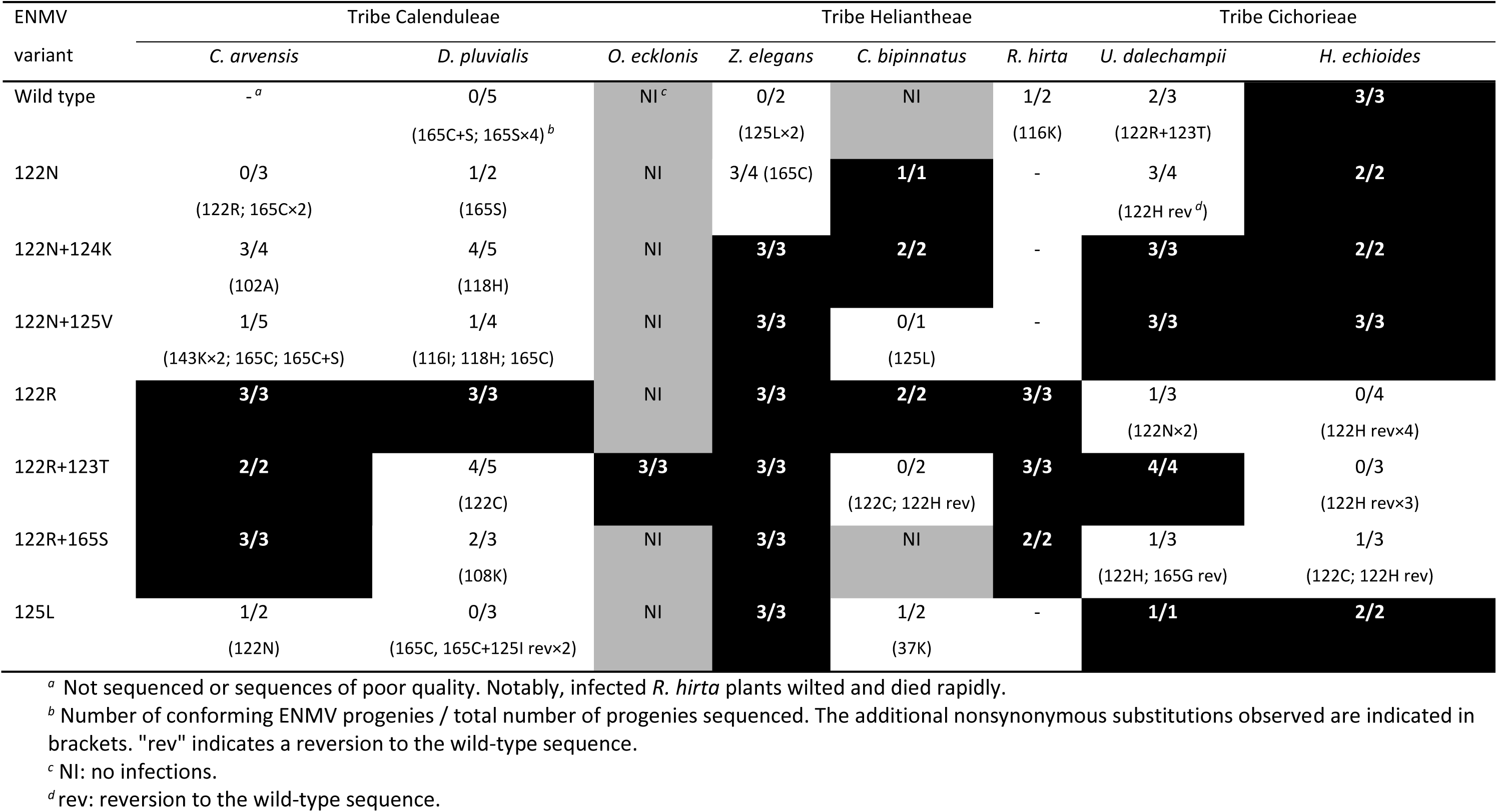
Sequence conformity of the VPg coding region of ENMV progenies in eight Asteraceae species with that of the original cDNA clones. The only *Calendula officinalis* plant infected with the 122R+123T mutant had a conforming VPg sequence. Cells in grey correspond to absence of infection. Cells in black correspond to situations where all progenies had a conforming VPg sequence.

In *C. arvensis*, the VPg sequence of the progeny of mutant viruses matched that of the corresponding cDNA clone for the three mutants containing the 122R mutation, which also corresponded to the mutants with the highest infection rates (77.5 to 85.0%) and the most frequent mutation detected in the ENMV populations evolved in that host (5 out of 8; Table 1). In the majority of cases (3/4), the VPg sequence of the progeny also matched that of the cDNA of the 122N + 124K double mutant, which also had a fairly high infection rate (65.0%). The progeny of the three remaining mutants (122N, 122N+125V, 125L) was consistent with the corresponding cDNA clone in up to 50% of cases. It should be noted that these three mutants were not detected in the experimentally evolved populations and had the lowest infection rates of the seven mutants (30.0 to 35.6%). These results indicate that three mutations or mutation combinations were sufficient to confer on ENMV a high level of adaptation to

*C. arvensis*, while the other four mutations or mutation combinations conferred an increase in adaptation compared with the wild-type virus and suggest that additional mutations are required for optimal adaptation, such as mutations 102A, 122N, 122R, 143K, 165C or 165S which were detected in viral progenies in infected *C. arvensis* plants (Table 3). It should be noted that these secondary mutations were located in the same positions (122 and 165) or regions (102 and 143) as those conferring adaptation to ENMV in *Z. elegans* or *C. arvensis*.

In *C. arvensis*, systemic viral load in infected plants followed the same trends as infection rates for all ENMV mutants examined. The highest loads were observed for the three mutants containing the 122R mutation, which also had the highest infection rates (Fig. 4A). In *Z. elegans*, the ranking of mutants according to systemic viral load varied according to experimental block (Figs. 4B and 4C). However, the mutants with extreme loads remained the same, with the 125L, 122R and 122N+125V mutants, which had among the highest loads, having significantly higher loads than the 122N mutant, the lowest, in both blocks. The 125L mutant with the highest loads was also the most frequently detected among experimentally-evolved populations (4 out of 8; Table 1).

**Figure 4.**
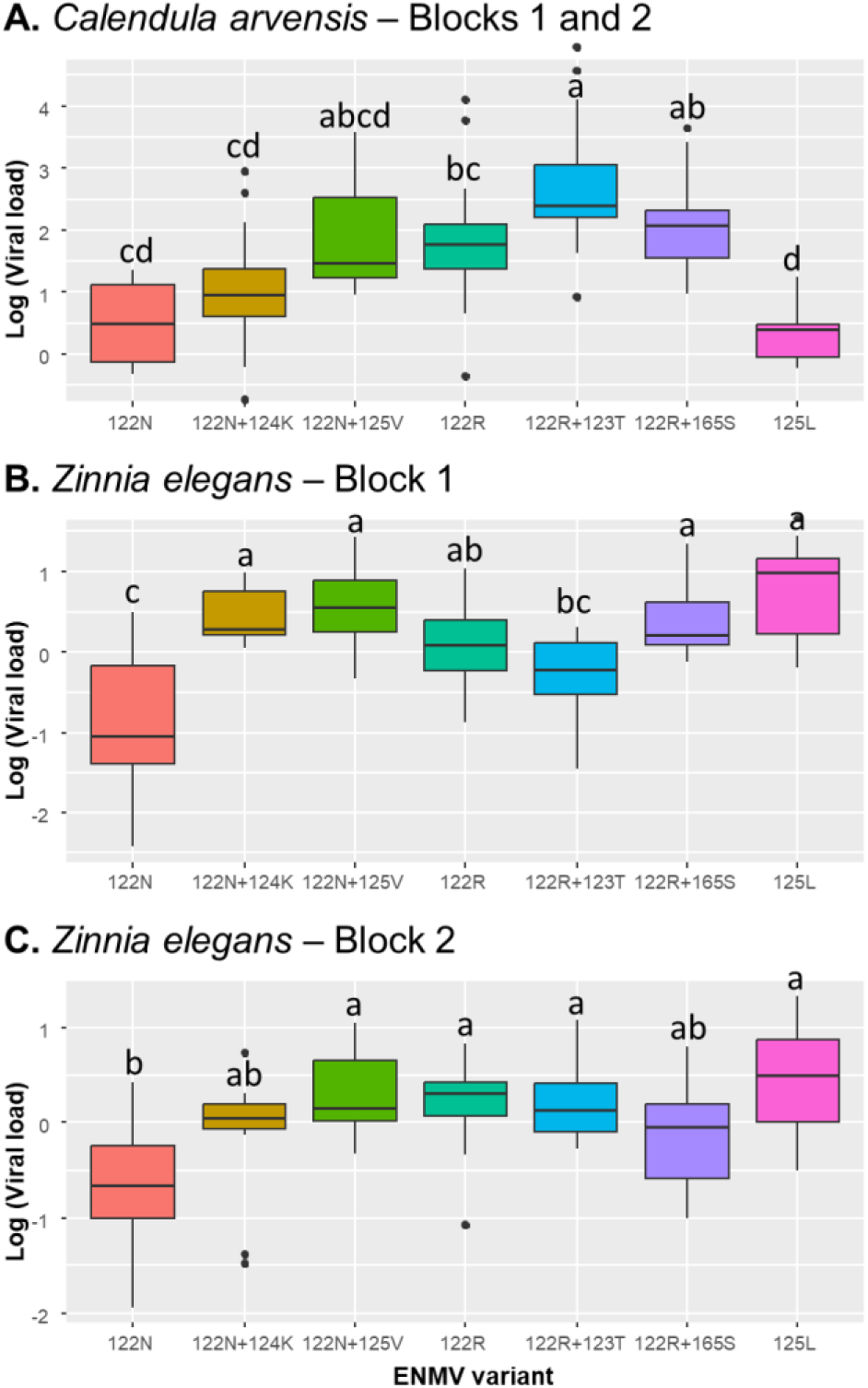
Systemic load of ENMV VPg mutants in *Calendula arvensis* (A) and *Zinnia elegans* (B, C). Relative viral loads were estimated by quantitative DAS-ELISA and are presented on a logarithmic scale for greater clarity. A significant interaction between the effect of the ENMV mutant and the effect of the experimental block was detected for *Z. elegans*, so separate analyses were performed for block 1 (B) and block 2 (C). Systemic viral loads in *C. arvensis* and *Z. elegans* were transformed using the Box-Cox method and the logarithmic function, respectively, prior to analysis of variance to satisfy normality of residuals and homogeneity of variances. Letters indicate significant differences between ENMV variants.

### Cross-adaptations of ENMV to Asteraceae species induced by VPg mutations are widespread and not related to host taxonomy

A striking result is the demonstration that the mutation pairs 122R+123T and 122R+165S, selected during experimental evolution in *C. arvensis*, confer a high level of adaptation in *Z. elegans* and, conversely, the 125L mutation and the mutation pairs 122N+124K and 122N+125V, selected in *Z. elegans*, confer a significant level of adaptation in *C. arvensis* (Tables 1 and 2). In addition, although it could not be used for reverse genetics analyses due to viral instability or low fitness, we noted that the 165C mutation alone was selected during experimental evolution in both *C. arvensis* and *Z. elegans.* These « cross-adaptation » effects, also known as cross-infectivity (Moury et al. 2014) or cross-virulence (Fournet et al. 2013), occur when a mutation (or more) selected in a parasite’s genome by one host (the primary host) also enhances adaptation to another host (the secondary host) that was not involved in the selection process. One hypothesis behind these cross-adaptations is that *C. arvensis* and *Z. elegans* are interchangeable in terms of ENMV evolution because they belong to two closely related tribes in the phylogeny of the Asteraceae family, namely Calenduleae and Heliantheae, respectively, whereas the likely species of origin of the ENMV isolate are *L. sativa* or *T. pratensis*, which, along with *C. endivia*, belong to the more distant tribe Cichorieae (Mandel et al. 2019).

To further test this hypothesis, we inoculated the wild-type ENMV clone and its seven VPg mutants into additional species belonging to these three tribes. In this experiment, whatever the ENMV variant, 100% infection was observed in *Cichorium endivia* (tribe Cichorieae). Among the 11 additional plant species tested, no infection was observed in *Catananche coerulea* (tribe Cichorieae) or in *Helianthus annuus*, *Echinacea purpurea* or *Coreopsis grandiflora* (tribe Heliantheae). Only one *Calendula officinalis* plant (tribe Calenduleae) was infected (with mutant 122R+123T). Results obtained with the other plant species are shown in Tables 2 and 3. In *Helminthotheca echioides* (tribe Cichorieae), the infection rate varied between 30 and 50% depending on the ENMV variant, with no significant differences. For the remaining five species, significant differences in infectivity were detected between variants. In *Osteospermum ecklonis* (tribe Calenduleae), a significantly higher infection rate was detected for the 122R+123T mutant (21.4%) than for the other variants, including the wild type (no infections). Similarly, some mutants displayed significantly higher infection rates than the wild type in *Cosmos bipinnatus* (tribe Heliantheae) (mutants 122R, 122R+123T and 122N+124K), in *Rudbeckia hirta* (tribe Heliantheae) (mutants 122R and 122R+123T) and in *Urospermum dalechampii* (tribe Cichorieae) (mutants 122N, 122N+124K, 122N+125V and 122R+123T). Finally, in *Dimorphotheca pluvialis* (tribe Calenduleae), the 125L mutant showed a significantly lower infection rate (32.5%) than the wild type (86.0%), with the other mutants showing no significant differences from the wild-type.

In these additional host species, the VPg sequence of mutant progenies matched the corresponding cDNA clone in a majority of cases (65%; 64/99) (Table 3). Second-site mutations affected positions involved in ENMV adaptation (122, 123, 125 or 165) in 84% of cases (32/38), suggesting adaptive roles. Several were reversions to the wild-type. Five of the remaining six mutations affected positions 108, 116 and 118, also within a critical VPg region for host adaptation (see below). A significant relationship was observed between infection rates and second-site mutations: mean infection was 32.6% when no progeny matched the clone sequence versus 60.4% when all did (p-value = 0.027; Kruskall-Wallis test).

Overall, VPg mutations (or pairs of mutations) induced cross-adaptations in 21.5% of cases (14/65) among Asteraceae species (Tables A and B in Supplementary Text S1). Contrary to expectations, these were not more frequent within the same tribe, nor between the closely related tribes Calenduleae and Heliantheae, compared between species of the more distant tribes Calenduleae and Cichorieae, or Heliantheae and Cichorieae (Tables A and C in Supplementary Text S1).

Altogether, these reverse genetics analyses demonstrated that nonsynonymous substitutions at codon positions 122, 123, 124, 125 and 165 of the VPg cistron, alone or in combination, were involved in ENMV adaptation to Asteraceae species, increasing its infection rate and, for some of them, their systemic viral load (Table 2 and Fig. 4).

## Discussion

### Capacity of EE to test propensity of host jumps

The EE carried out revealed rapid adaptation, i.e. in three to four infection cycles corresponding to three or four months of evolution, of a genetically-homogeneous ENMV strain to two new host species: the wild plant *C. arvensis* and the ornamental plant *Z. elegans*. In the other three host species (*L. sativa*, *T. pratensis* and *C. endivia*), we found no evidence of phenotypic evolution of ENMV in terms of infection rate, systemic viral load or virulence. This could be due to the fact that ENMV is already well adapted to these hosts, since isolate ENMV-FR was collected in a plot of *L. sativa* and ENMV shows a high prevalence in *T. pratensis* in the vicinity of the sampled plot.

Evidence for adaptation of ENMV to *C. arvensis* and *Z. elegans* is supported by two main observations. First, infection rates increased markedly over successive passages: from 22.4% to 62.5% in *C. arvensis* and from 37.5% to 95.8% in *Z. elegans* within the first two to three infection cycles, eventually reaching 100% in both hosts. Second, systemic viral loads were higher for the final evolved populations than for either the ancestral strain or ENMV lineages maintained in non-selective hosts (*L. sativa*, *C. endivia*, and *T. pratensis*), as demonstrated by the cross-inoculation assay. This evidence is further supported by the observation of parallel fixation of nonsynonymous substitutions in the VPg cistron of independent ENMV populations evolved in these two hosts and the demonstration that these substitutions (or pairs of these substitutions) confer a clear gain in adaptation to ENMV in terms of infection rates. Altogether, these results indicate that ENMV has the capacity to adapt to new host species through a short evolutionary time and a small number of genetic changes.

The infection rate and viral accumulation results provide also strong evidence for reciprocal cross-adaptation of ENMV between *C. arvensis* and *Z. elegans*. With regard to viral accumulation, in *C. arvensis*, ENMV populations evolved in *Z. elegans* accumulated about threefold higher than the populations evolved in *L. sativa* or *T. pratensis*. Conversely, in *Z. elegans*, ENMV populations evolved in *C. arvensis* reached accumulation levels about 20-fold higher than the ancestral strain and populations evolved in *L. sativa*, *C. endivia*, or *T. pratensis*.

These results are consistent with the local adaptation signal of ENMV detected in *Z. elegans* and *C. arvensis* (Supplementary Fig. S3). These adaptations are associated with only minor costs in alternative host species, detected solely in *C. endivia*. In this host, populations evolved in *C. arvensis* and *Z. elegans* exhibited lower infection rates and systemic loads, respectively, compared with the other evolved populations (Fig. 3A,B). These fitness costs may underlie the local-versus-foreign adaptation pattern observed in this species (Supplementary Fig. S3).

### Similarity of potyvirus evolutionary mechanisms involved in adaptation to new host species and eIF4E (eukaryotic initiation factor 4E)-mediated immunity at host species level

The adaptation of plant viruses to new host species has not been extensively studied, unlike their adaptation to new plant genotypes of the same species. In particular, little is known about the number of mutations required for adaptation, the dynamics of their accumulation in viral genomes, the involvement of recombination and the importance of pre-adaptation in host jumps (Holmes and Drummond 2007). We might have assumed that the greater the genetic distance between the virus primary and secondary hosts, the greater the number of mutations (or evolutionary steps) required for adaptation. Our results on ENMV indicate that as few mutations are required for viral adaptation to new species as has been observed for viral adaptation to new plant genotypes, i.e. one or two mutations in most cases (Moury et al. 2021). However, this observation may be biased, as only the most rapid host adaptations and the simplest viral genetic determinants may have been studied experimentally and published, regardless of the genetic distance between hosts.

It should also be noted that only the very first stage of viral host jump has been studied here, namely the evolution of a genetic compatibility between the virus and the new host species. This genetic compatibility does not guarantee that the virus will be efficiently transmitted from one plant to another in the new host species, and that it will persist over time in this host population.

Analysis of a database on the host range of plant viruses has shown that host barriers are much more frequent and/or impermeable when plant species belong to different botanical families than when they belong to the same family. Our results are consistent with this observation, since we observed a rapid adaptation of ENMV to new species and genera of Asteraceae. A previous analysis of ENMV host range also showed that 15/15 species tested in the Asteraceae were infected, while 0/9 species outside the Asteraceae were infected, after experimental inoculation (Desbiez et al. 2017).

We showed that nonsynonymous substitutions at codon positions 122, 123, 124, 125 and 165 of the VPg cistron determined ENMV adaptation to different Asteraceae species. Comparisons between single and double mutants suggest additive effects in some cases, explaining the frequent selection of multiple mutants. For instance, 122N+124K and 122R+123T often outperformed 122N and 122R, respectively, in some hosts (Tables 1 and 2). However, additional testing of single mutants (e.g., 123T, 124K) is needed to confirm true additivity. These effects were not universal, and epistatic interactions likely occur, as some double mutants performed worse than single ones, for example, 122R+165S failed to infect *C. bipinnatus*, whereas 122R alone infected 47% of plants (Table 2).

The progeny of these viruses in different plant species showed additional second-site nonsynonymous mutations at codon positions 37, 102, 108, 116, 118, 122, 143 and 165 (Table 3), some of which may also be involved in ENMV adaptation. VPg positions 102, 116, 118, 122 and 165 coincide with VPg positions involved in resistance breakdown (i.e. adaptation to new genotypes of the same plant species) for other potyviruses (PVY, TVMV - *Potyvirus nicotianavenamaculae* -, TuMV and MWMV - *Potyvirus citrullimoroccense* -) (Fig. 5). A few studies have also shown the role of the VPg in potyvirus adaptation to new host species. Rajamäki and Valkonen (1999) showed that substitutions at positions corresponding to codon positions 118 and 187 of the VPg cistron of ENMV conferred to potato isolates of PVA an adaptation to *Nicandra physalodes*, another *Solanaceae*. Again, the first of these positions belongs to the critical region for resistance breakdown. Similarly, a mutation at the codon position corresponding to ENMV position 167 conferred to TEV an adaptation to *A. thaliana* (Agudelo-Romero et al. 2008) and mutations at codon positions corresponding to ENMV positions 116 and 165 conferred to PPV an adaptation to *A. thaliana* and *C. foetidum* (Martinez-Turiño et al. 2021). These positions coincide or are close to the adaptive mutations of ENMV. Several other positions in ENMV VPg (108, 123, 124, 125) that revealed mutations during our experiments (Table 3) are located within or close to the critical region for resistance breakdown or adaptation to new host species, which is located in the center of the VPg (extending from position 102 to 122 in ENMV and from 101 to 121 in PVY; Ayme et al 2007).

**Figure 5.**
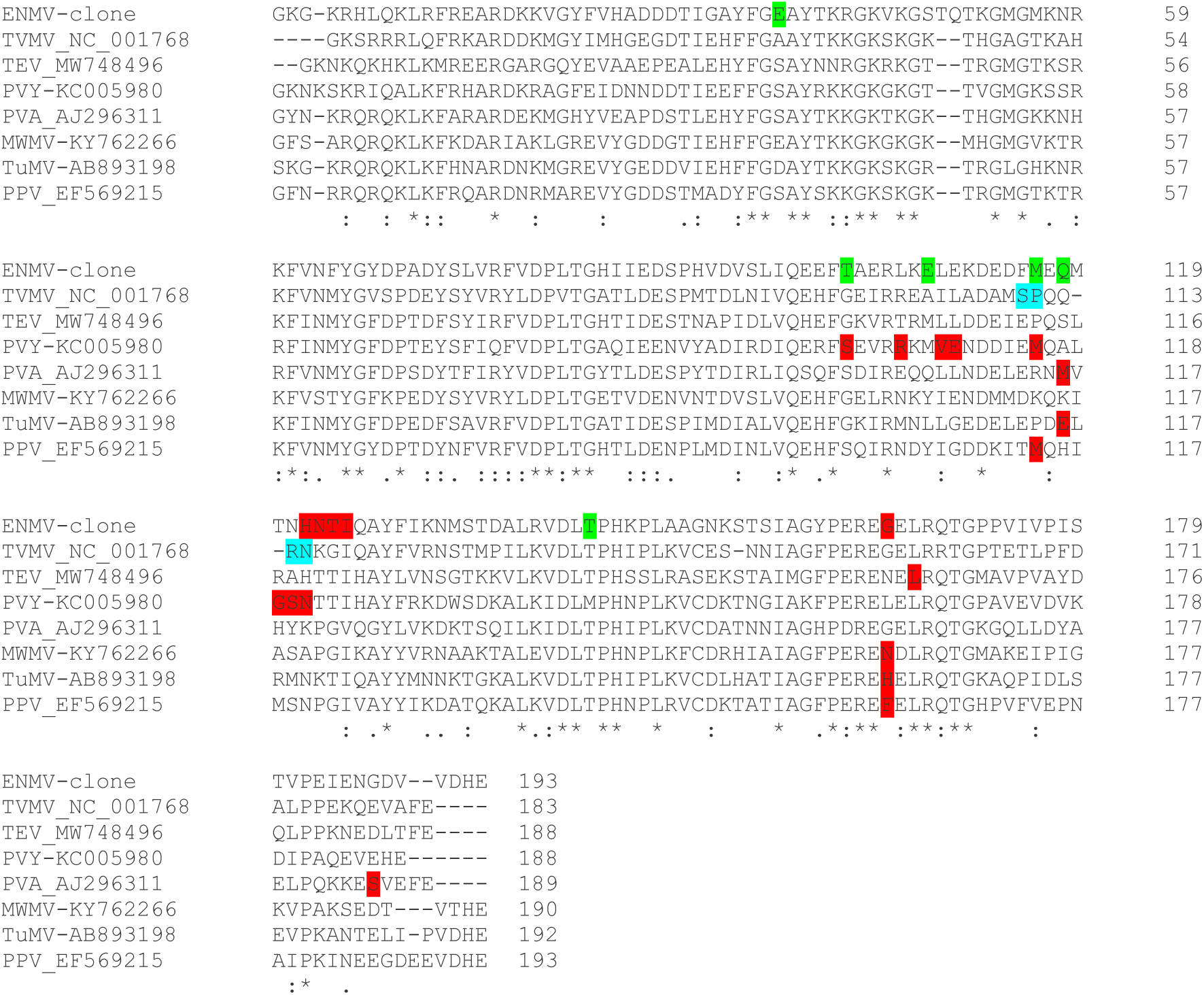
Alignment of VPg amino acid sequences of ENMV and other potyviruses showing key positions involved in adaptation to the plant host. Data are from Nicolas et al. (1997) for tobacco vein mottling virus (TVMV), Agudelo-Romero et al. (2008) for tobacco etch virus (TEV), Ayme et al. (2006, 2007) and Janzac et al. (2014) for potato virus Y (PVY), Rajamäki and Valkonen (1999) for potato virus A (PVA), Pechar et al. (2022) for Moroccan watermelon mosaic virus (MWMV), Gallois et al. (2010) for turnip mosaic virus (TuMV) and Martínez-Turiño et al. (2021) for plum pox virus (PPV). Positions underlined in red have been shown to be individually involved in host adaptation. Positions underlined in blue were shown to be involved in host adaptation as a whole but not individually. Positions underlined in green were observed in ENMV progenies during infection of different plant species, but were not shown to be individually involved in host adaptation.

n conclusion, these results suggest that, in potyviruses, the breakdown of resistance in certain plant genotypes of a given plant species operates by the same mechanisms as adaptation to new plant species (i.e. a small number of nonsynonymous substitutions in either a small central region of the VPg cistron or at particular positions at its C-terminal extremity, corresponding to codon positions 165 or 167 of ENMV VPg cistron).

For host plants of the other potyviruses PVY, TVMV, TuMV and MWMV, the broken down resistances correspond to eIF4E-mediated plant immunity, and VPg mutations confer resistance breakdown either by restoring VPg binding to the mutated eIF4E protein or by allowing its binding to another member of the small eIF4E family (Gallois et al. 2018). It is therefore possible that the adaptation of ENMV to new species of Asteraceae, determined by VPg mutations, also takes place through hijacking of eIF4E proteins by the virus in these plant species.

### Mutations with pleiotropic (cross-adaptation) effects in terms of host range are frequent in ENMV VPg and unlinked with plant taxonomy

One of the most striking findings was the high frequency of cross-adaptation effects caused by pleiotropic effects in the ENMV VPg cistron (21.5%; Table A in Supplementary Text S1). These cross-adaptation effects were not linked to the host plant taxonomy: the primary and secondary hosts did not belong to the same tribe within the Asteraceae more frequently than expected by chance, nor did they belong to the closely-related tribes Calenduleae and Heliantheae more frequently than expected by chance (Tables B and C in Supplementary Text S1). An explanation for the observed cross-adaptation effects may lie in the distribution of adaptive mutations within highly restricted regions of the viral VPg protein and the host eIF4E protein, whereas the remainder of these proteins is subject to strong functional constraints. Such a structural organization increases the likelihood that mutations arising in these limited regions will have pleiotropic effects, as reported for ENMV and other potyviruses (Moury et al., 2014; Martínez-Turiño et al., 2021). Adaptive changes in VPg and eIF4E appear to involve subtle modifications that alter their molecular interactions. Notably, these effects can be driven by a small number of selected mutations located in specific protein regions. Importantly, these mutations neither reflect nor disrupt the broader taxonomic relationships among viruses (in the case of VPg) or among host plants (in the case of eIF4E). This functional decoupling from phylogenetic relatedness may explain why the cross-adaptation patterns observed in this study are not associated with host plant taxonomy.

## Conclusion

This study demonstrates the usefulness and efficiency of combining experimental evolution and cross-inoculation assays to uncover virus host shifts and identify the mutations underlying adaptation. In a companion paper, Roques et al. (2026) inferred from the same set of cross-inoculation data, a multi-host phenotypic fitness landscape for ENMV using a Fisher’s geometrical model framework in which each host is represented as a fitness peak in a shared phenotypic space, and where infection success is linked to this geometry through an explicit mechanistic model combining three categories of infection of the new host, namely through the direct establishment of the predominant variant of the viral population, through the establishment of a minor variant already present in the inoculum (standing variation) or through the establishment of a variant that emerged de novo after inoculation, under a strong-selection weak-mutation regime. Beyond estimating distances among host-specific phenotypic optima, including that of the ancestral strain, this approach also quantified host permissiveness (through the width of the fitness peak) and target-specific infection efficiency (i.e. how efficiently phenotypic suitability translates into successful infection), thereby separating geometric constraints on host shifts from host-dependent effects on establishment. The inferred landscape reveals two clearly separated clusters: one including the three Cichorieae hosts *Cichorium endivia*, *Lactuca sativa* and *Tragopogon pratensis*, and a second including *Calendula arvensis* and *Zinnia elegans*. In addition, the inferred permissiveness parameters indicate that *Lactuca sativa* and *Tragopogon pratensis* are highly permissive hosts, whereas *Cichorium endivia* is more intermediate and *Calendula arvensis* and *Zinnia elegans* behave as more restrictive hosts. This is consistent with the stronger molecular signature of VPg evolution observed during adaptation to these restrictive hosts in the experimental study, as well as with the limited evidence for fitness costs associated with these adaptations in more permissive hosts. Overall, the experimental and modelling approaches are highly complementary: the former identifies the mutations and phenotypes underlying host adaptation, whereas the latter provides a parsimonious and mechanistically interpretable framework for understanding how host community structure shapes the emergence potential of ENMV and for discussing possible springboard effects.

## Acknowledgements

This study was performed thanks to the INRAE experimental facilities (IEPV) of the Plant Pathology unit (https://eng-pathologie-vegetale.paca.hub.inrae.fr/infrastructures/prophyle/experimental-facilities) and the LBM platform of INRAE PACA. Our warmest thanks go to Joël Béraud, Nathalie Truglio, Michel Pascal and Jérémy Théodore of the IEPV for plant maintenance and to Judith Hirsch, Karine Nozeran, Sylvain Piry, Loup Rimbaud, Alexandra Schoeny and Eric Verdin for assistance with the experimental evolution and cross-inoculation assays. We also thank Brigitte Maisonneuve for providing the lettuce seeds and Guillaume Martin for interesting discussions about the project.

## Abbreviations

ANOVA: analysis of variance
CTV: citrus tristeza virus
DAS-ELISA: double-antibody sandwich enzyme-linked immunosorbent assay
dpi: day post inoculation
EE: experimental evolution
ENMV: endive necrotic mosaic virus
ESS: effective sample size
LMV: lettuce mosaic virus
LOO: leave one out
MWMV: Moroccan watermelon mosaic virus
ns: not significant
PPV: plum pox virus
PRSV: papaya ringspot virus
PVA: potato virus A
PVY: potato virus Y
RYMV: rice yellow mottle virus
TEV: tobacco etch virus
TuMV: turnip mosaic virus
TVMV: tobacco vein mottling virus
VPg: viral protein genome-linked.

## References

Agudelo-Romero P, Carbonell P, Perez-Amador MA, Elena SF (2008) Virus adaptation by manipulation of host’s gene expression. PLoS ONE, 3, e2397. 10.1371/journal.pone.0002397

Ayme V, Souche S, Caranta C, Jacquemond M, Chadœuf J, Palloix A, Moury B (2006) Different mutations in the genome-linked protein VPg of potato virus Y confer virulence on the *pvr2^3^* resistance in pepper. Molecular Plant-Microbe Interactions, 19, 557–563. 10.1094/MPMI-19-0557

Ayme V, Petit-Pierre J, Souche S, Palloix A, Moury B (2007) Molecular dissection of the potato virus Y VPg virulence factor reveals complex adaptations to the *pvr2* resistance allelic series in pepper. Journal of General Virology, 88,1594–1601. 10.1099/vir.0.82702-0

Belabess Z, Dallot S, El-Montaser S, Granier M, Majde M, Tahiri A, Blenzar A, Urbino C, Peterschmitt M (2015) Monitoring the dynamics of emergence of a non-canonical recombinant of *Tomato yellow leaf curl virus* and displacement of its parental viruses in tomato. Virology, 486, 291–306. 10.1016/j.virol.2015.09.011

Bürkner P-C (2017) brms: An R package for Bayesian multilevel models using Stan. Journal of Statistical Software, 80, 1–28. 10.18637/jss.v080.i01

Bürkner P-C (2018) Advanced Bayesian multilevel modeling with the R package brms. The R Journal, 10, 395 411. 10.32614/RJ-2018-017

Carpenter B, Gelman A, Hoffman MD, Lee D, Goodrich B, Betancourt M, Brubaker M, Guo J, Li P, Riddell A (2017) Stan: A probabilistic programming language. Journal of Statistical Software, 76, 1–32. 10.18637/jss.v076.i01

Chen KC, Chiang CH, Raja JA, Liu FL, Tai CH, Yeh SD (2008) A single amino acid of NIaPro of *Papaya ringspot virus* determines host specificity for infection of papaya. Molecular Plant-Microbe Interactions, 21, 1046–1057. 10.1094/mpmi-21-8-1046

Desbiez C, Joannon B, Wipf-Scheibel C, Chandeysson C, Lecoq H (2009) Emergence of new strains of *Watermelon mosaic virus* in South-Eastern France: evidence for limited spread but rapid local population shift. Virus Research, 141, 201–208. 10.1016/j.virusres.2008.08.018

Desbiez C, Chandeysson C, Lecoq H, Moury B (2012) A simple, rapid and efficient way to obtain infectious clones of potyviruses. Journal of Virological Methods, 183, 94–97. 10.1016/j.jviromet.2012.03.035

Desbiez C, Chandeysson C, Lecoq H (2014) A short motif in the N-terminal part of the coat protein is a host-specific determinant of systemic infectivity for two potyviruses. Molecular Plant Pathology, 15, 217–221. 10.1111/mpp.12076

Desbiez C, Schoeny A, Maisonneuve B, Berthier K, Bornard I, Chandeysson C, Fabre F, Girardot G, Gognalons P, Lecoq H, Lot H, Millot P, Nozeran K, Simon V, Tepfer M, Verdin E, Wipf-Scheibel C, Moury B (2017) Molecular and biological characterization of two potyviruses infecting lettuce in southeastern France. Plant Pathology, 66, 970–979. 10.1111/ppa.12651

Fournet S, Kerlan MC, Renault L, Dantec JP, Rouaux C, Montarry J (2013) Selection of nematodes by resistant plants has implications for local adaptation and cross-virulence. Plant Pathology, 62, 184–193. 10.1111/j.1365-3059.2012.02617.x

Gal-On A, Meiri E, Elman C, Gray D, Gaba V (1997) Simple hand-held devices for the efficient infection of plants with viral-encoding constructs by particle bombardment. Journal of Virological Methods, 64, 103–110. 10.1016/s0166-0934(96)02146-5

Gallois JL, Charron C, Sánchez F, Pagny G, Houvenaghel MC, Moretti A, Ponz F, Revers F, Caranta C, German-Retana S (2010) Single amino acid changes in the turnip mosaic virus viral genome-linked protein (VPg) confer virulence towards *Arabidopsis thaliana* mutants knocked out for eukaryotic initiation factors eIF(iso)4E and elF(iso)4G. Journal of General Virology, 91, 288–293. 10.1099/vir.0.015321-0

Gallois JL, Moury B, German-Retana S (2018) Role of the genetic background in resistance to plant viruses. International Journal of Molecular Sciences, 19, 2856. doi:10.3390/ijms19102856

Gelman A, Rubin DB (1992) Inference from iterative simulation using multiple sequences. Statistical Science, 7, 457–472. 10.1214/ss/1177011136.

Gómez P, Rodríguez-Hernández A, Moury B, Aranda M (2009) Genetic resistance for the sustainable control of plant virus diseases: breeding, mechanisms and durability. European Journal of Plant Pathology, 125, 1–22. 10.1007/s10658-009-9468-5

Holmes EC, Drummond A (2007) The evolutionary genetics of viral emergence. Current Topics in Microbiology and Immunology 315, 51–66. 10.1007/978-3-540-70962-6_3

Janzac B, Tribodet M, Lacroix C, Moury B, Verrier JL, Jacquot E (2014) Evolutionary pathways to break down the resistance of allelic versions of the PVY resistance gene *va*. Plant Disease, 98, 1521–1529. 10.1094/pdis-11-13-1126-re

Johnston CK, Waterhouse T, Wiens M, Mondick J, French J, Gillespie WR (2024) Bayesian estimation in NONMEM. CPT: Pharmacometrics & Systems Pharmacology, 13, 192–207. 10.1002/psp4.13088

Kawecki TJ, Ebert D (2004) Conceptual issues in local adaptation. Ecology Letters, 7, 1225–1241. 10.1111/j.1461-0248.2004.00684.x

Mandel JR, Dikow RB, Siniscalchi CM, Thapa R, Watson LE, Funk VA (2019) A fully resolved backbone phylogeny reveals numerous dispersals and explosive diversifications throughout the history of Asteraceae. Proceedings of the National Academy of Sciences of the United States of America, 116, 14083–14088. 10.1073/pnas.1903871116

Martínez-Turiño S, Calvo M, Bedoya LC, Zhao M, García JA (2021) Virus host jumping can be boosted by adaptation to a bridge plant species. Microorganisms, 9, 805. 10.3390/microorganisms9040805

Moury B, Janzac B, Ruellan Y, Simon V, Ben Khalifa M, Fakhfakh H, Fabre F, Palloix A (2014) Interaction patterns between potato virus Y and eIF4E-mediated recessive resistance in the *Solanaceae*. Journal of Virology, 88, 9799–9807. 10.1128/jvi.00930-14

Moury B, Desbiez C (2020) Host range evolution of potyviruses: A global phylogenetic analysis. Viruses, 12, 111. doi:10.3390/v12010111

Moury B, Tamisier L, Fabre F, Rimbaud L (2021) Peut-on prédire la durabilité des résistancesLe cas des virus. In: Lannou C, Roby D, Ravigné V, Hannachi M, Moury B, L’immunité des plantes, Quae, Versailles, France, pp. 297–307.

Navaud O, Barbacci A, Taylor A, Clarkson JP, Raffaele S (2018) Shifts in diversification rates and host jump frequencies shaped the diversity of host range among *Sclerotiniaceae* fungal plant pathogens. Molecular Ecology, 27, 1309–1323. 10.1111/mec.14523

Nicolas O, Dunnington SW, Gotow LF, Pirone TP, Hellmann GM (1997) Variations in the VPg protein allow a potyvirus to overcome *va* gene resistance in tobacco. Virology, 237, 452–459. 10.1006/viro.1997.8780

Panstruga R, Moscou MJ (2020) What is the molecular basis of nonhost resistance? Molecular Plant-Microbe Interactions, 33, 1253–1264. 10.1094/mpmi-06-20-0161-cr

Pechar GS, Donaire L, Gosalvez B, García-Almodovar C, Sánchez-Pina MA, Truniger V, Aranda MA (2022) Editing melon eIF4E associates with virus resistance and male sterility. Plant Biotechnology Journal, 20, 2006–2022. 10.1111/pbi.13885

Poulicard N, Pinel-Galzi A, Traoré O, Vignols F, Ghesquière A, Konate G, Hébrard E, Fargette D (2012) Historical contingencies modulate the adaptability of *Rice yellow mottle virus*. PLoS Pathogens, 8, e1002482. 10.1371/journal.ppat.1002482

Rajamäki ML, Valkonen JPT (1999) The 6K2 protein and the VPg of potato virus A are determinants of systemic infection in *Nicandra physaloides*. Molecular Plant-Microbe Interactions, 12, 1074–1081. 10.1094/mpmi.1999.12.12.1074

Romay G, Lecoq H, Desbiez C (2015) *Melon chlorotic mosaic virus* and associated alphasatellite from Venezuela: genetic variation and sap transmission of a begomovirus-satellite complex. Plant Pathology, 64, 1224–1234. 10.1111/ppa.12342

Roques L, Papaïx J, Martin G, Forien R, Lenormand T, Soubeyrand S, Berthier K, Moury B (2026) Inferring the multi-host fitness landscape of endive necrotic mosaic virus from cross-inoculation experiments. bioRxiv, 10.64898/2026.03.18.712764

Suehiro N, Natsuaki T, Watanabe T, Okuda S (2004) An important determinant of the ability of *Turnip mosaic virus* to infect *Brassica* spp. and/or *Raphanus sativus* is in its P3 protein. Journal of General Virology, 85, 2087–2098. 10.1099/vir.0.79825-0

Svanella-Dumas L, Verdin E, Faure C, German-Retana S, Gognalons P, Danet JL, Marais A, Candresse T. (2014) Adaptation of *Lettuce mosaic virus* to *Catharanthus roseus* Involves mutations in the central domain of the VPg. Molecular Plant-Microbe Interactions, 27, 491–497. 10.1094/mpmi-10-13-0320-r

Tamisier L, Lacombe S, Caranta C, Gallois JL, Moury B (2023) Virus evolution faced to multiple host targets: The potyvirus-pepper case study. Current Topics in Microbiology and Immunology, 439, 121–138. 10.1007/978-3-031-15640-3_3

Tamura K, Stecher G, Peterson D, Filipski A, Kumar S (2013) MEGA6: Molecular Evolutionary Genetics Analysis Version 6.0. Molecular Biology and Evolution, 30, 2725–2729. 10.1093/molbev/mst197

Tatineni S, Robertson CJ, Garnsey SM, Dawson WO (2011) A plant virus evolved by acquiring multiple nonconserved genes to extend its host range. Proceedings of the National Academy of Sciences of the United States of America, 108, 17366–17371. 10.1073/pnas.1113227108

Thines M (2019) An evolutionary framework for host shifts – jumping ships for survival. The New Phytologist, 224, 605–617. 10.1111/nph.16092

Truniger V, Nieto C, González-Ibeas D, Aranda M (2008) Mechanism of plant eIF4E-mediated resistance against a *Carmovirus* (Tombusviridae): cap-independent translation of a viral RNA controlled in cis by an (a)virulence determinant. The Plant Journal, 56, 716–727. 10.1111/j.1365-313x.2008.03630.x

Vassilakos N, Simon V, Tzima A, Johansen E, Moury B (2016) Genetic determinism and evolutionary reconstruction of a host jump in a plant virus. Molecular Biology and Evolution, 33, 541–553. 10.1093/molbev/msv222

Vehtari A, Gelman A, Gabry J (2017) Practical Bayesian model evaluation using leave-one-out cross-validation and WAIC. Statistics and Computing, 27, 1413–1432. 10.1007/s11222-016-9696-4

Wallis CM, Stone AL, Sherman DJ, Damsteegt VD, Gildow FE, Schneider WL (2007) Adaptation of plum pox virus to a herbaceous host (*Pisum sativum*) following serial passages. Journal of General Virology, 88, 2839–2845. 10.1099/vir.0.82814-0

Woolhouse MEJ, Gowtage-Sequeria S (2005) Host range and emerging and reemerging pathogens. Emerging Infectious Diseases, 11, 1842–1847. 10.3201/eid1112.050997

