## Supplementary Text S1 for "Common molecular determinants underlie potyvirus host species jumps and resistance breakdown"

**Supplementary Text S1. Analysis of cross-adaptations of ENMV between Asteraceae species.**

**Table A.** Summary of cross-adaptation data based on Table 2. « Yes » : significantly higher infection rate than the wild-type ENMV. « No » : infection rate not significantly higher than that of the wild-type ENMV. « Not relevant » : in these situations, the primary and secondary hosts correspond to the same species ; therefore, these are not cases of cross-adaptations.

| Variant | Primary host for adaptation | Secondary host for cross adaptation (Calenduleae) | | | Secondary host for cross adaptation (Heliantheae) | | | | | | Secondary host for cross adaptation (Cichorieae) | | | Total |
|---|---|---|---|---|---|---|---|---|---|---|---|---|---|---|
| | | *C. arvensis* | *C. officinalis* | *O. ecklonis* | *C. grandiflora* | *C. bipinnatus* | *E. purpurea* | *H. annuus* | *R. hirta* | *Z. elegans* | *C. caerulea* | *H. echioides* | *U. dalechampii* | |
| 122R+123T | *C. arvensis* | Not relevant | No | Yes | No | Yes | No | No | Yes | Yes | No | No | Yes | 5/11 |
| 122R+165S | *C. arvensis* | Not relevant | No | No | No | No | No | No | No | Yes | No | No | No | 1/11 |
| 122R | *C. arvensis* + *Z. elegans* | Not relevant | No | No | No | Yes | No | No | Yes | Not relevant | No | No | No | 2/10 |
| 122N+125V | *Z. elegans* | Yes | No | No | No | No | No | No | No | Not relevant | No | No | Yes | 2/11 |
| 122N | *Z. elegans* | Yes | No | No | No | No | No | No | No | Not relevant | No | No | Yes | 2/11 |
| 122N+124K | *Z. elegans* | Yes | No | No | No | Yes | No | No | No | Not relevant | No | No | Yes | 3/11 |
| 125L | *Z. elegans* | Yes | No | No | No | No | No | No | No | Not relevant | No | No | No | 1/11 |
| Total | | | | | | | | | | | | | | 14/65 |

**Table B.** Contingency table corresponding to the test of the null hypothesis : ENMV cross-adaptations are equally probable within and between Asteraceae tribes. This hypothesis is not rejected according to Fisher's exact test (p-value = 0.25). The ENMV 122N mutant was excluded because of its redundance with the 122N+125V mutant (Table A above).

| | Gains of adaptation | No adaptation gain |
|---|---|---|
| Within tribe | 4 (18.2%) | 22 (81.8%) |
| Between tribes | 10 (25.6%) | 29 (74.4%) |

**Table C.** Contingency table corresponding to the test of the null hypothesis : ENMV cross-adaptations are equally likely (i) within Asteraceae tribes or between Calenduleae and Heliantheae tribes and (ii) between Cichorieae and another tribe. This hypothesis is not rejected according to Fisher's exact test (p-value = 0.41). The 122N ENMV mutant was excluded because of its redundance with the 122N+125V mutant (Table A above).

| | Gains of adaptation | No adaptation gain |
|---|---|---|
| Within tribe or between Calenduleae and Heliantheae | 11 (23.4%) | 36 (76.6%) |
| Between Cichorieae and another tribe | 3 (16.7%) | 15 (83.3%) |
