## Supplementary figures and images for "Common molecular determinants underlie potyvirus host species jumps and resistance breakdown"

### Supplementary Figure S1

**Supplementary Figure S1. Strategy used to build the cDNA clones of ENMV VPg mutants.**

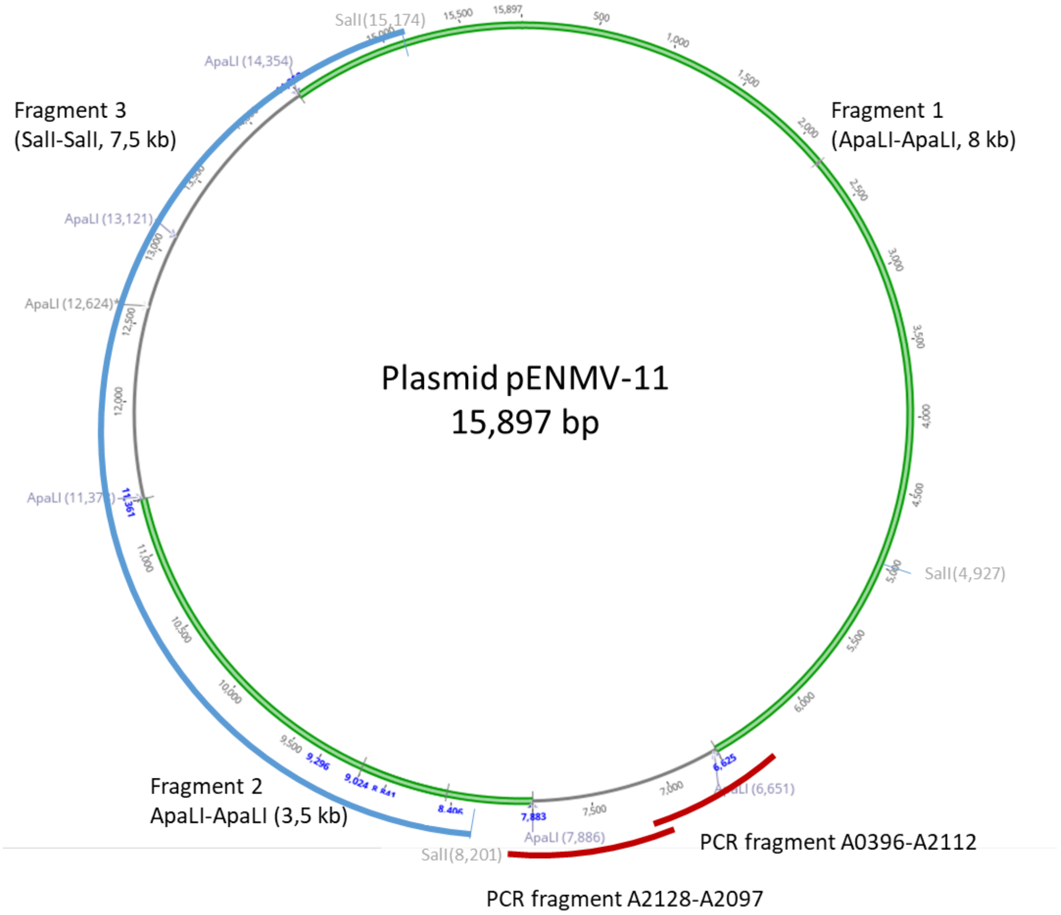
