## Supplementary Figure S2 for "Common molecular determinants underlie potyvirus host species jumps and resistance breakdown"

**Supplementary Figure S2. Predicted probability of infection by viral population type–plant species combination.** Tiles show predicted infection probabilities from a Bayesian binomial regression model (brms) including random effects for block and strain : lineage. The horizontal axis represents viral population types (initial strain 7098MP1 or populations evolved in the five different plant species), and the vertical axis represents the test plant species. Tile color indicates predicted probability (blue = low, yellow = high). The plot asymmetry shows strong interactions between viral population type and test plant species.

CH: *Cichorium endivia*; LA: *Lactuca sativa*; SA: *Tragopogon pratensis*; ZI: *Zinnia elegans*; SO: *Calendula arvensis*.

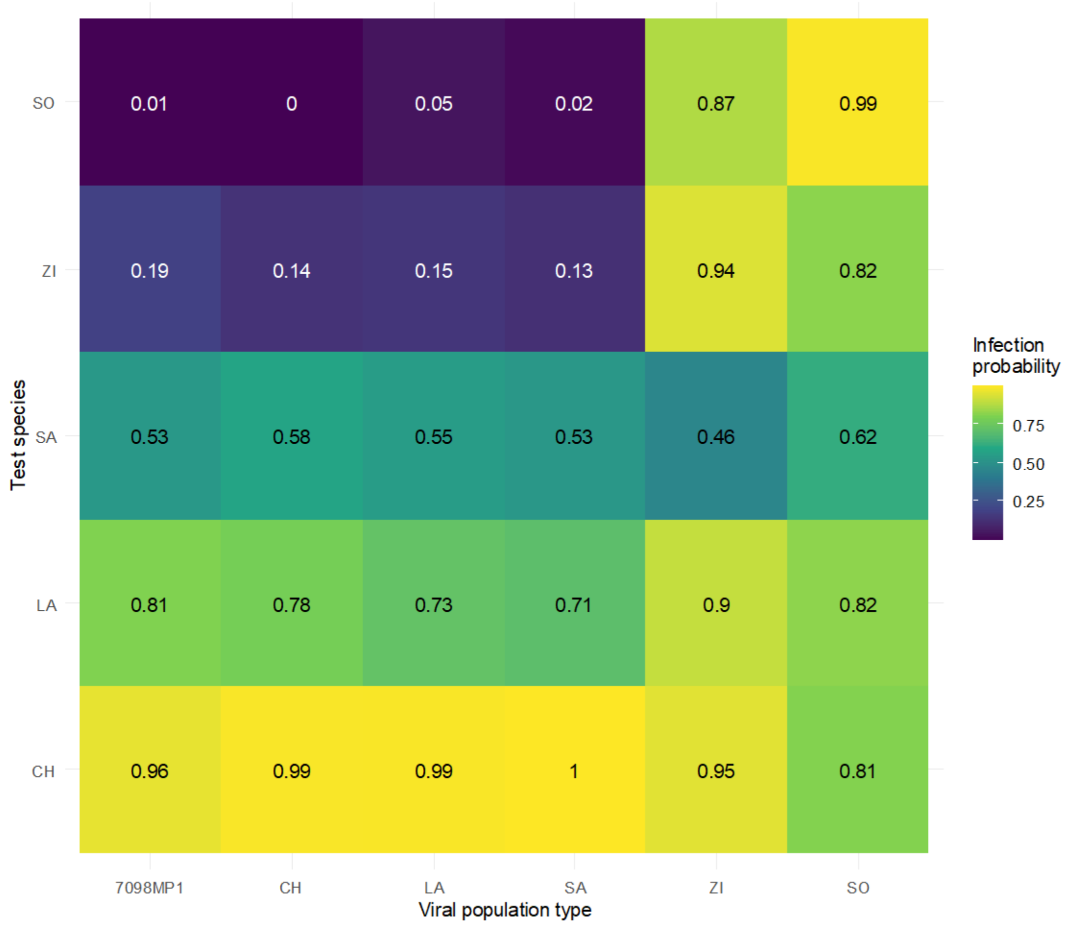
