## Supplementary Figure S3 for "Common molecular determinants underlie potyvirus host species jumps and resistance breakdown"

**Supplementary Figure S3. Analysis of local adaptation of ENMV to its plant host species after experimental evolution during six serial infection cycles, as assessed using the foreign-versus-local and home-versus-away frameworks.** Difference in infection probability between "Local" and "Foreign" (**A**) and between "Home" and "Away" (**B**) conditions for each host species (respectively, each viral population type). Points represent posterior mean differences (Local − Foreign or Home − Away), and error bars indicate 95% credible intervals from the Bayesian binomial mixed-effects model. Difference in viral load (log scale) between "Local" and "Foreign" (**C**) and between "Home" and "Away" (**D**) conditions for each host species (respectively, each viral population type). Points represent estimated contrasts (Local − Foreign or Home − Away) from the linear mixed-effects model, and error bars indicate 95% confidence intervals. The dashed horizontal lines indicate no difference between conditions.

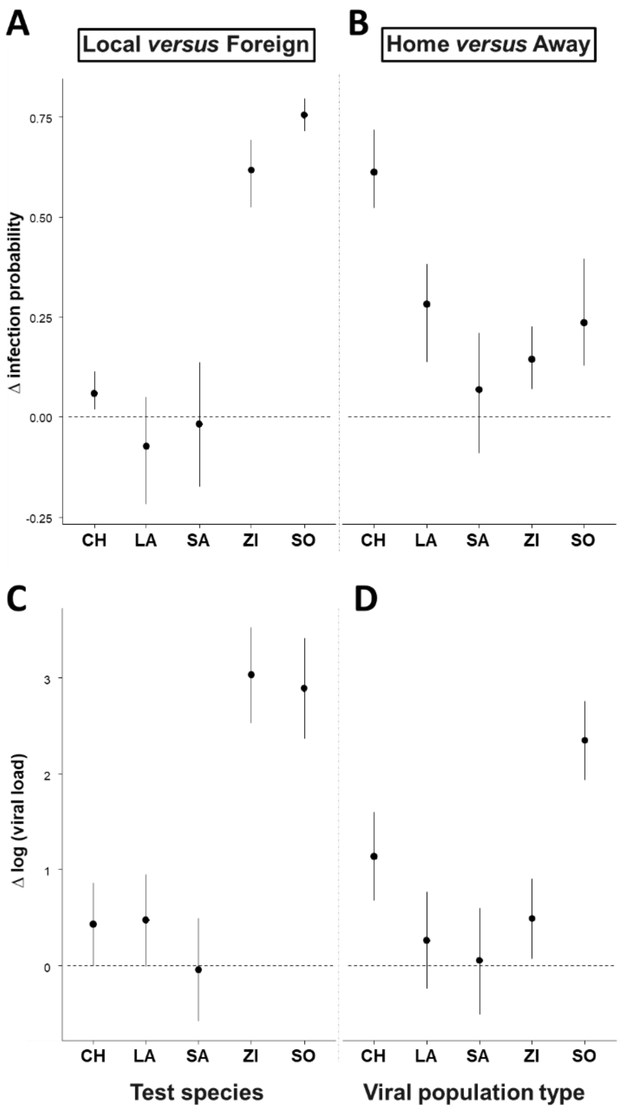
