## Supplementary Table S1 for "Common molecular determinants underlie potyvirus host species jumps and resistance breakdown"

**Supplementary Table S1. Primers used to obtain cDNA clones of VPg mutants of pENMV-11.**

| Primer name | Orientation[a] | Sequence[b] | Position in ENMV genome[c] | Coding region |
|---|---|---|---|---|
| A0396 | FW | CACACACTCGAGAACATCGC | 5857-5876 | CI |
| A2112 | RV | CCTTTTTGCAATGAGCGGGC | 6750-6769 | Pro |
| A2128 | FW | GACGTAGTTGATCACGAAGC | 6733-6752 | VPg-Pro |
| A2097 | RV | AAGTCTACACTTTCCTTGGAC | 7564-7583 | NIb |
| A2129 | FW | GAGCAAATGACAAAT**A**ACAACACAATCCA | 6520-6548 | VPg |
| A2130 | RV | TGGATTGTGTTGT**T**ATTTGTCATTTGCTC | 6520-6548 | VPg |

[a]: FW : forward ; RV : reverse.

[b]: bold and underlined : site-directed mutagenesis to introduce the H122N mutation.

[c]: based on GenBank accession number KU941946.
