## Supplementary Table S2 for "Common molecular determinants underlie potyvirus host species jumps and resistance breakdown"

**Supplementary Table S2. Versions of the R packages used.**

| Package | Version |
|---|---|
| abind | 1.4-8 |
| arrayhelpers | 1.1-0 |
| backports | 1.5.0 |
| bayesplot | 1.15.0 |
| boot | 1.3-30 |
| bridgesampling | 1.2-1 |
| brms | 2.23.0 |
| Brobdingnag | 1.2-9 |
| broom | 1.0.12 |
| broom.mixed | 0.2.9.7 |
| cachem | 1.1.0 |
| car | 3.1-5 |
| carData | 3.0-6 |
| cellranger | 1.1.0 |
| checkmate | 2.3.4 |
| class | 7.3-22 |
| cli | 3.6.5 |
| coda | 0.19-4.1 |
| codetools | 0.2-20 |
| cowplot | 1.2.0 |
| data.table | 1.18.2.1 |
| DescTools | 0.99.60 |
| devtools | 2.4.6 |
| DHARMa | 0.4.7 |
| digest | 0.6.39 |
| distributional | 0.7.0 |
| dplyr | 1.2.0 |
| e1071 | 1.7-17 |
| ellipsis | 0.3.2 |
| emmeans | 1.11.0 |
| estimability | 1.5.1 |
| evaluate | 1.0.5 |
| Exact | 3.3 |
| expm | 1.0-0 |
| farver | 2.1.2 |
| fastmap | 1.2.0 |
| forcats | 1.0.1 |
| Formula | 1.2-5 |
| fs | 1.6.6 |
| furrr | 0.3.1 |
| future | 1.69.0 |
| generics | 0.1.4 |

| | |
|---|---|
| ggdist | 3.3.3 |
| ggplot2 | 4.0.2 |
| ggpubr | 0.6.2 |
| ggsignif | 0.6.4 |
| gld | 2.6.8 |
| globals | 0.19.0 |
| glue | 1.8.0 |
| gridExtra | 2.3 |
| gtable | 0.3.6 |
| haven | 2.5.5 |
| hms | 1.1.4 |
| htmltools | 0.5.9 |
| httr | 1.4.7 |
| knitr | 1.51 |
| lattice | 0.22-6 |
| lifecycle | 1.0.5 |
| listenv | 0.10.0 |
| lme4 | 1.1-37 |
| lmerTest | 3.2-0 |
| lmom | 3.2 |
| loo | 2.9.0 |
| lubridate | 1.9.5 |
| magrittr | 2.0.4 |
| MASS | 7.3-60.2 |
| Matrix | 1.7-0 |
| matrixStats | 1.5.0 |
| memoise | 2.0.1 |
| minqa | 1.2.8 |
| multcomp | 1.4-29 |
| multcompView | 0.1-10 |
| mvtnorm | 1.3-3 |
| nlme | 3.1-168 |
| nloptr | 2.2.1 |
| numDeriv | 2016.8-1.1 |
| otel | 0.2.0 |
| parallelly | 1.46.1 |
| patchwork | 1.3.2 |
| pillar | 1.11.1 |
| pkgbuild | 1.4.8 |
| pkgconfig | 2.0.3 |
| pkgload | 1.5.0 |
| plyr | 1.8.9 |
| posterior | 1.6.1 |
| proxy | 0.4-29 |
| purrr | 1.2.1 |

| | |
|---|---|
| R6 | 2.6.1 |
| rbibutils | 2.3 |
| RColorBrewer | 1.1-3 |
| Rcpp | 1.1.0 |
| RcppParallel | 5.1.11-1 |
| Rdpack | 2.6.4 |
| readr | 2.1.6 |
| readxl | 1.4.5 |
| reformulas | 0.4.4 |
| remotes | 2.5.0 |
| reshape2 | 1.4.5 |
| rlang | 1.1.7 |
| rlist | 0.4.6.2 |
| rmarkdown | 2.30 |
| rootSolve | 1.8.2.4 |
| rstantools | 2.6.0 |
| rstatix | 0.7.3 |
| rstudioapi | 0.18.0 |
| S7 | 0.2.1 |
| sandwich | 3.1-1 |
| scales | 1.4.0 |
| sessioninfo | 1.2.3 |
| stringi | 1.8.7 |
| stringr | 1.6.0 |
| survival | 3.6-4 |
| svUnit | 1.0.8 |
| tensorA | 0.36.2.1 |
| TH.data | 1.1-5 |
| tibble | 3.3.1 |
| tidybayes | 3.0.7 |
| tidyr | 1.3.2 |
| tidyselect | 1.2.1 |
| tidyverse | 2.0.0 |
| timechange | 0.4.0 |
| tzdb | 0.5.0 |
| usethis | 3.2.1 |
| vctrs | 0.7.1 |
| withr | 3.0.2 |
| xfun | 0.56 |
| xtable | 1.8-4 |
| yaml | 2.3.12 |
| zoo | 1.8-14 |
